# Calibrated analysis framework for nanopore direct RNA sequencing uncovers cell-specific m⁶A proportions at conserved sites

**DOI:** 10.1101/2025.11.02.686099

**Authors:** Denise Ohnezeit, Elene Loliashvili, Gregory Putzel, Ruth Verstraten, Adam T. Silhavy, Jianheng Liu, Luke S. Nicholson, Alejandro Pironti, Samie R. Jaffrey, Daniel P. Depledge, Angus C. Wilson

## Abstract

Nanopore direct RNA sequencing (DRS) coupled with Dorado modification-aware basecalling enables mapping of epitranscriptomic modifications including N^6^-methyladenosine (m^6^A) at the level of individual RNAs. However, the sensitivity, specificity, and reproducibility of this method remain unclear and have only recently begun to be addressed through systematic benchmarking studies. Here, we aimed to establish a best-practice workflow for DRS-based epitranscriptomic analyses. Specifically, we evaluated multiple Dorado versions and models using RNA isolated from primary cells and unmodified *in vitro* transcribed RNAs. We further utilized an m^6^A methyltransferase inhibitor as a specificity control. We established that stringent filtering is necessary to reduce the contribution of background and found that Dorado predictions captured an increasing proportion of GLORI sites detected at high m^6^A/A proportions. Further, by applying DRS to a human primary fibroblast and a HD10.6 neuron dataset, we detected differences in the predicted m^6^A/A proportions at conserved sites between these datasets. Our study thus presents the first systematic comparison of Dorado and GLORI from the same input RNA and expands characterization of the m^6^A epitranscriptome to fibroblasts and neurons.

## Introduction

Epitranscriptomics describes the occurrence of chemical modifications on RNA that regulate stability, localization, splicing, translation, or decay (Gilbert and Nachtergaele 2023). To date, over 170 RNA modifications are known to occur across all classes of RNA, including eukaryotic mRNA (Cappannini et al. 2023). Of these, at least thirteen different chemical modifications have been characterized in mRNAs, comprising both cap-proximal and internal modifications (Anreiter et al. 2021). By contrast, current evidence indicates that lncRNAs harbor a more limited repertoire of modifications, primarily N6-methyladenosine (m⁶A), 5-methylcytosine (m⁵C), and N7-methylguanosine (m⁷G) (Yang et al. 2024). One of the best-studied and most abundant internal mRNA modifications is m^6^A. Functional studies of m^6^A methylases, demethylases, and m^6^A binding proteins, as well as sequencing-based mapping methods, have increased our understanding of m^6^A as a reversible and dynamic modification that fine-tunes nearly all aspects of mRNA metabolism (Zaccara et al. 2019; He and He 2021; Murakami and Jaffrey 2022; Gilbert and Nachtergaele 2023; Sendinc and Shi 2023).

In addition to m⁶A, other widespread and functionally important marks include m⁵C, which is less abundant but influences mRNA export and stability, pseudouridine (Ψ), which contributes to mRNA stability, translation and splicing, and adenosine-to-inosine (A to I) editing, which is involved in splicing, stability, and recoding events that contribute to protein diversity (Sun et al. 2023).

Techniques based on short-read (Illumina) sequencing have advanced in recent years to achieve single-nucleotide resolution and quantitative mapping of RNA modifications. The approaches that have emerged to map m^6^A can be broadly classified as antibody-based, enzyme-assisted or chemical conversion methods (Moshitch-Moshkovitz et al. 2024).

Early antibody-based techniques (Dominissini et al. 2012; Meyer et al. 2012; Linder et al. 2015) provided the first transcriptome-wide insights into m^6^A architecture, albeit at limited resolution. Enzyme-assisted methods (Meyer 2019; Xiao et al. 2023) have improved precision but can introduce enzymatic biases. These biases are avoided with GLORI-Sequencing (GLORI), which selectively deaminates unmethylated adenosines, enabling theoretically unbiased, quantitative, single-nucleotide mapping of m⁶A (Liu et al. 2023).

Despite advances, the broad utility of these approaches is limited by a lack of isoform specificity, as well as biases associated with amplification, specificity of antibodies or variable adenosine conversion rates (Diensthuber and Novoa 2025). The short-read length inherent to these approaches also limits the ability to determine whether multiple modifications occur on the same individual RNA molecule. Oxford Nanopore Technologies (ONT) direct RNA sequencing (DRS) offers an alternative to these methods by analyzing native RNA molecules that have retained their modifications (Garalde et al. 2018). This enables modification detection in an isoform-resolved, single-molecule context and, in principle, allows examination of the co-occurrence of multiple modifications along individual RNA molecules. As single RNAs pass through nanopores, signal changes are interpreted to determine canonical and modified residues at single-nucleotide resolution on individual transcripts (Jain et al. 2016; Viehweger et al. 2019). For the now discontinued DRS SQK-RNA002 chemistry used with R9 flowcells, multiple research groups developed algorithms for detecting RNA modifications. These methods can be divided into error rate-based and current signal-based (Abebe et al. 2022). While the recent release of the DRS SQK-RNA004 chemistry and RNA-specific flowcells reduced the utility of many of these tools, introduction of a modification-aware basecaller (Dorado) provides an all-in-one toolkit for basecalling, RNA modification detection, and poly(A) tail length estimation. An early RNA modification-aware Dorado model (v3.0.1) supported m^6^A detection exclusively within the DRACH context, while a later model (v5.0.0) introduced detection of Ψ and all-context m^6^A. A more recent model (v5.1.0) supports both DRACH-context and all-context m^6^A detection, together with Ψ, inosine, and m^5^C (github.com/nanoporetech/Dorado).

In a recent overview of this emerging technology, Cruciani and Novoa highlighted the importance of rigorous evaluation of Dorado’s limitations and advocated for the establishment of best practices to ensure consistency and accuracy going forward (Cruciani and Novoa 2026). Furthermore, Diensthuber and Novoa emphasized the need for direct comparisons between Dorado and orthogonal approaches such as GLORI using matched input material, as most benchmarking studies rely on previously published datasets rather than side-by-side experimental comparisons (Diensthuber and Novoa 2025).

To date, several studies have evaluated Dorado in a benchmarking context. Two studies compared Dorado with m6Anet using SQK-RNA004 chemistry and reported improved performance of Dorado, while also incorporating IVT controls to assess false-positive predictions (Zou et al. 2025; Ghohabi Esfahani et al. 2026). Incorporating selective METTL3 inhibition as a complementary, modification-specific negative control could further strengthen the assessment of m^6^A detection specificity. Both studies also applied predicted m^6^A/A proportion-based filtering to define high-confidence Dorado modification sites, using thresholds of 10% in Zou et al. and 20% in Ghobabi Esfahani et al.; however, the effect of the chosen predicted m^6^A/A proportion threshold on site detection and false-positive reduction was not systematically evaluated. Consequently, a comprehensive set of best-practice parameters for Dorado-based modification detection has yet to be established. Notably, although Zou et al. compared Dorado-derived predictions with GLORI-identified sites, the GLORI data were obtained from previously published datasets rather than generated from matched input material, leaving direct comparisons between Dorado and GLORI on the same RNA samples unaddressed (Cruciani et al. 2025; Diensthuber and Novoa 2025; Zou et al. 2025; Ghohabi Esfahani et al. 2026). By directly comparing m^6^A detection by Dorado and GLORI on the same RNA inputs, our study overcomes this limitation. Here we systematically benchmark Dorado using poly(A) RNA from primary normal human dermal fibroblasts (NHDFs) and functional human neurons produced through terminal differentiation of HD10.6 cells (Raymon et al. 1999), along with IVT RNA. We included treatment with STM2457, an enzymatic inhibitor of the primary m^6^A methyltransferase METTL3 (Yankova et al. 2021), and compared the modification-detection performance of multiple Dorado models released since the tool’s introduction. While our analysis is centered on the sensitivity and specificity of m^6^A detection in DRS datasets, we also benchmark the ability of Dorado to reliably detect m^5^C, Ψ, and inosine. Together, our data provide a robust workflow for DRS-based modification detection that reliably predicts transcriptome-wide, isoform-level, site-specific proportions of RNA modifications. By expanding m^6^A mapping on primary human fibroblasts and functional human neurons, we uncover mostly conserved m^6^A profiles but reveal increased predicted m^6^A/A proportions in neuronal cells.

## Results

### Systematic benchmarking of ONT’s Dorado within human fibroblast and neuronal datasets

To evaluate the performance of ONT’s basecalling tool Dorado across different human cell types, we selected primary NHDFs and differentiated HD10.6 cells as a neuron-like model. Prior to isolation of total RNA, cells were treated for 48h with a small-molecule inhibitor of METTL3 (STM2457) or with vehicle (DMSO). We ensured that this treatment had no effect on cell viability by performing a CellTiter Glo® assay (**Fig. S1**). We then isolated poly(A) RNA and subjected the same input material to both DRS and GLORI. The same material was also used to generate IVT RNA (**Fig. 1A**). Dorado basecalling using software version v.0.9.0 and basecalling model version v.5.1.0 resulted in over 13M reads using poly(A) RNA from NHDFs treated with DMSO or STM2457, while we obtained over 5M and 10M reads from DMSO-and STM2457-treated HD10.6 cells, respectively. The rate of primary alignment to the human genome (hg38) was over 98% for all the sequenced samples. A detailed overview of read statistics generated by DRS can be found in **Supplementary Table 1**.

**Figure 1.**
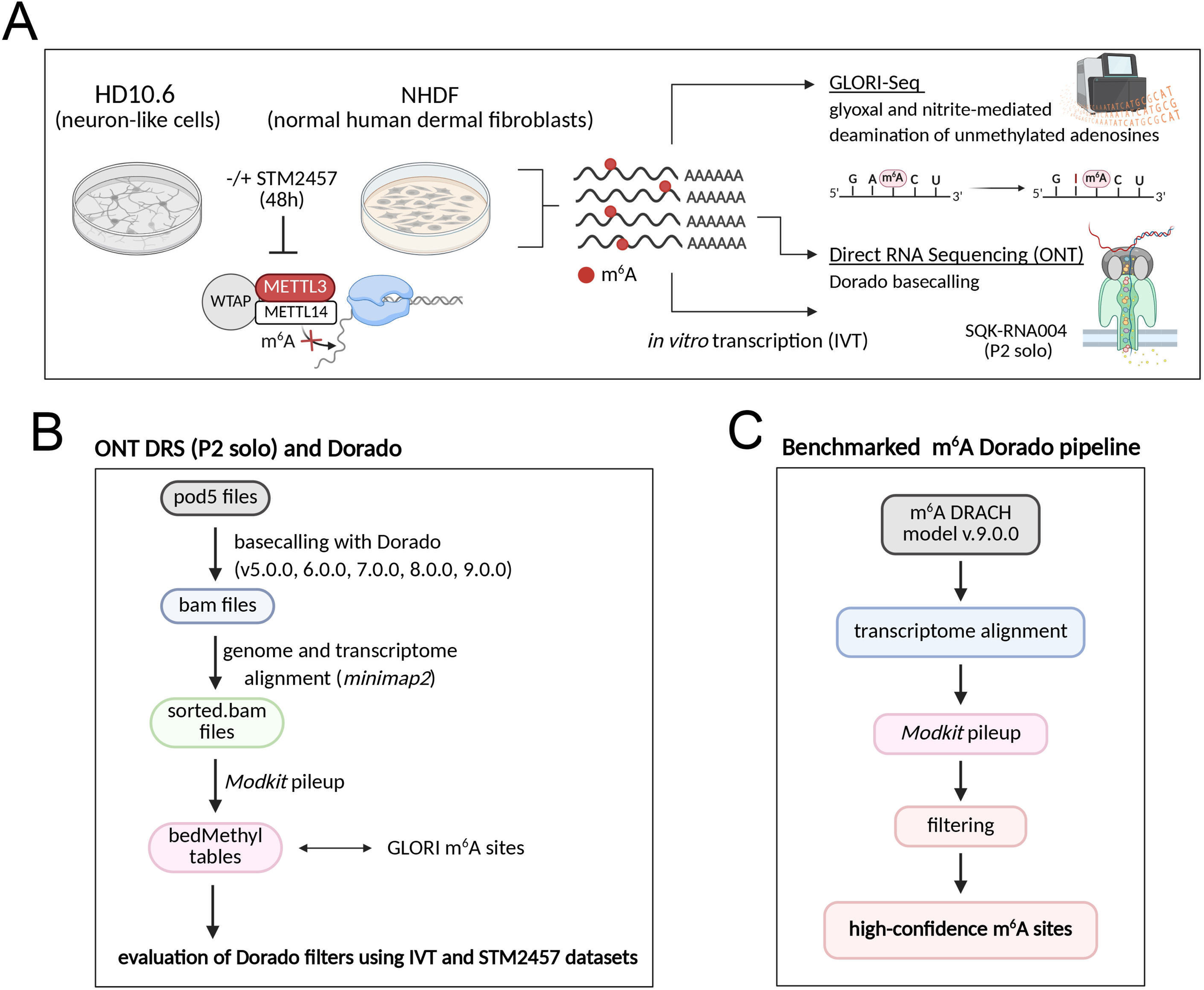
Workflow for m^6^A mapping in different human cell types using orthogonal sequencing approaches. **(A)** Schematic of the experimental setup. RNA was isolated from NHDFs and differentiated HD10.6 cells treated with STM2457 or DMSO for 48h. Poly(A) RNA was subjected to DRS and GLORI from the same input material. IVT RNA was generated in parallel to assess modification calls arising from background signal. **(B)** Workflow for benchmarking Dorado basecalling. Reads were processed with Dorado versions v.0.5.0 through v0.9.0, aligned to the human genome or transcriptome, and analyzed with ONT’s Modkit tool. Multiple filtering strategies were tested against IVT controls and METTL3 inhibition (STM2457). Resulting m⁶A sites were compared with GLORI data from the same input RNA. **(C)** Final benchmarked Dorado pipeline to detect high confidence m^6^A sites.

To systematically benchmark Dorado, we performed basecalling using software versions v0.5.0 through v0.9.0. Reads were aligned to the human genome and transcriptome to obtain comprehensive information on isoform-level modification detection. Further, we explored different aspects of detected modifications using the outputs generated by ONT’s Modkit tool. We assessed multiple filtering strategies using base and modification probabilities, as well as predicted modification proportion distributions, and evaluated them based on the IVT data and sensitivity to METTL3 inhibition. By filtering out background signal using multiple control layers, we determined high-confidence m^6^A sites (**Fig. 1B**). Finally, we compared these high-confidence predictions with sites identified by GLORI performed on the same input RNA. Using these results, we developed a benchmarked pipeline that filters m^6^A sites predicted by Dorado to minimize background signal detection without impacting sensitivity (**Fig. 1C**).

### Stringent filtering of modification probabilities removes false positives

During modification-aware basecalling, each nucleotide is assigned a basecall probability score ranging from 0 to 1. Similarly, individual modified nucleotides are assigned a modification probability score in the same range. To establish a baseline for background signal, we compared basecall and m^6^A modification probability distributions between a human genome-aligned IVT poly(A) RNA dataset and DMSO-and STM2457-treated NHDF datasets, all processed using Dorado v0.9.0 (**Fig. 2A,B**, **Fig. S2**). Basecall probabilities for adenosine and m^6^A modification probabilities from both the all-context and DRACH-context models were grouped into probability bins for comparison.

**Figure 2.**
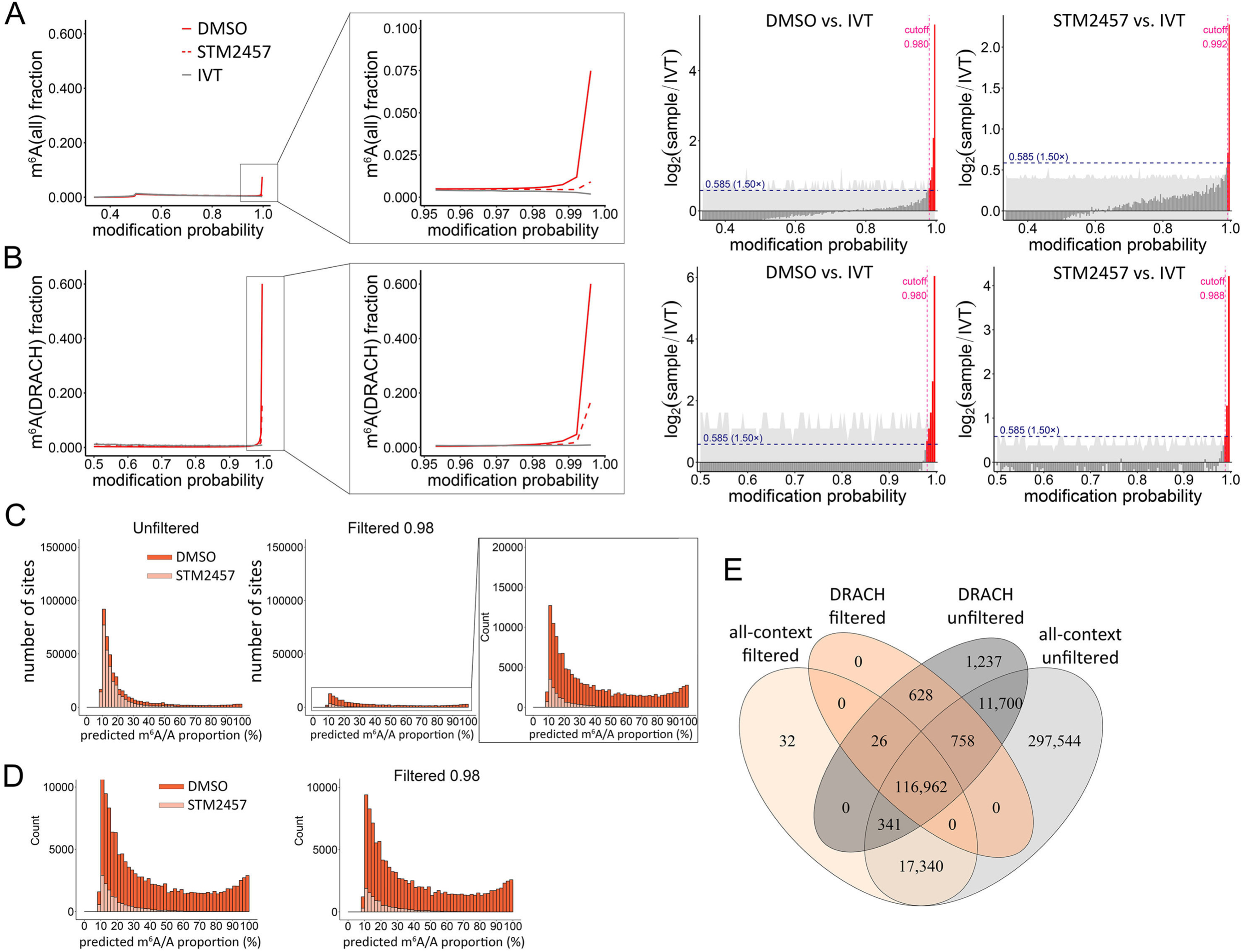
Modification probability of 0.98 and predicted m^6^A/A proportion score of ≥ 10% are necessary cutoffs to identify m^6^A sites accurately. **(A)** (left) Dorado v0.9.0-generated modification probability distributions for all-context m^6^A from genome-aligned DMSO-and STM2457-treated NHDFs plotted against the distributions obtained from the IVT NHDF poly(A) RNA. DMSO probability score distribution is shown as a red solid line, STM2457 – red dashed line, IVT – grey solid line. (right) Bar plots showing the per-bin log2(sample/IVT) ratio of modification probability fractions for the DMSO-and STM2457-treated NHDFs conditions relative to the unmodified IVT reference. The horizontal navy dashed line marks the fold-enrichment threshold used to call enrichment. Bars are coloured by significance relative to the fold-enrichment threshold (red, log2 ≥ 1.5-fold; dark grey, not significant). The light grey ribbon indicates the 95% bootstrap confidence interval around the log2 ratio estimate at each bin. The vertical pink dashed line indicates the derived modification probability cutoff above which sites are considered genuinely modified. **(B)** Same as in (A), but for DRACH-context m^6^A sites. **(C)** Predicted m^6^A/A proportion (≥ 10%) distributions for all-context m^6^A from DMSO-(in red) or STM2457-treated (in pink) NHDF datasets (genome-aligned) are plotted using a 0.98 modification probability cutoff and a filter for ≥ 20 reads. **(D)** Same as in (C), but for DRACH-context m^6^A sites. **(E)** Overlap of m^6^A sites between all-context and DRACH models (A basecall accuracy ≥ 80%, coverage ≥ 20 reads, predicted m^6^A/A proportion ≥ 10%), comparing non-filtered to filtered (modification probability > 0.98) outputs.

We observed that 75-90% of A basecalls (in all datasets) have probability scores above 0.8 (**Fig. S2A**). Since all A bases are considered during all-context m^6^A calling, the fraction of m^6^A sites is substantially smaller when compared to DRACH-context m^6^A sites (**Fig. 2A,B**, **Fig. S2B,C**). The modification probability distributions are effectively identical for DMSO-treated, STM2457-treated, and IVT datasets below 0.98, while the profiles differ substantially at probabilities ≥ 0.98. This suggests that sites detected below 0.98 are likely background signal, indicating that only high-probability m^6^A calls can be considered reliable. Consistent with this interpretation, METTL3 inhibition with STM2457 markedly reduced the number of detected m^6^A sites with modification probabilities ≥ 0.98, compared with DMSO-treated samples, in both the all-context and DRACH-only datasets (**Fig. 2A,B**, **Fig. S2B,C**).

To identify specific modification probability cutoffs for each comparison between biological samples and the IVT dataset, we established a statistical framework in which modification probabilities were divided into bins, and enrichment of modification calls in the biological sample relative to IVT was expressed as the log2-transformed sample/IVT ratio. The Pruned Exact Linear Time (PELT) algorithm was then used to identify the point at which this ratio shifted from the baseline distribution, indicating divergence of the biological signal from the IVT background. The modification probability cutoff was subsequently defined as the first bin following the detected change point that exceeded the specified fold-enrichment threshold. This analysis independently supported the use of stringent modification probability cutoffs to distinguish modification-associated signal from the IVT background (**Fig. 2A,B**).

To assess the contribution of background signal to modification detection across different modification probability thresholds (0.5, 0.7, 0.9, 0.98, and 0.99), 200,000 reads were randomly subsampled from each dataset in ten iterations, and modification probability distributions were determined for each subsample. Background signal was estimated by comparing modification counts in biological samples with those detected in the corresponding IVT controls at each threshold. In NHDF samples, the m^6^A DRACH sites showed a relatively low contribution of background signal (∼10%) even at a probability threshold of 0.5, which decreased further at more stringent thresholds (**Fig. S3A**). By contrast, the m^6^A all-context model showed substantially higher background signal (∼35% at ≥0.5), decreasing to <5% at the highest thresholds. Analysis of discrete modification probability ranges showed a similar trend (**Fig. S3B**), with background signal decreasing significantly at higher modification probabilities. Statistical significance was assessed using pairwise comparisons derived from linear mixed-effects models, with sample included as a random effect and Benjamini-Hochberg correction for multiple comparisons. The reduction in background signal in all-context m^6^A detection was less pronounced following STM2457 treatment, indicating that background signal becomes proportionally more prominent as the number of genuine modifications decreases. Similar patterns were observed in HD10.6 cells (**Fig. S4A,B**).

Furthermore, we examined modification probability distributions for Ψ, inosine and m^5^C, each of which can be basecalled with Dorado v0.9.0 (**Fig. S5**). For Ψ and I, the modification probability distributions for the STM2457 and DMSO datasets only diverge from the IVT dataset at ≥ 0.99 modification probability, supporting the use of this stringent threshold to reduce the impact of background signal (**Fig. S5A,B**). Consistent with this, the estimated background signal decreased with increasing modification probability, although they remained comparatively high (∼20-30%) even at stringent thresholds (**Fig. S5A,B, Fig. S6A, Fig. S7A**). Analysis across discrete modification probability intervals showed the same trend, with progressively lower error rates at higher probabilities (**Fig. S5A,B, Fig. S6B, Fig. S7B**). By contrast, m^5^C modification probability distributions were similar between all three datasets, and no filter could be adequately established (**Fig. S5C**). While this likely reflects a low rate of m^5^C installation, it also indicates that the m^5^C model utilized by Dorado requires further refinement.

Modification probability distributions have similar profiles for transcriptome-aligned NHDF data (**Fig. S8**) as well as genome-and transcriptome-aligned data from HD10.6 cells (**Fig. S9, Fig. S10**). Of note, the fraction of high-probability inosine sites is increased in STM2457-treated cells, compared to DMSO-treated ones in genome-aligned data (**Fig. S5B, Fig. S9D**), which points to increased numbers of retained introns in the METTL3-inhibited sample. Accordingly, this effect is absent in transcriptome-aligned data (**Fig. S8D, Fig. S10D**).

While IVT transcriptomes serve as excellent negative controls, we also considered whether the RNA calibration strand (RCS) that is added during DRS library preparation could serve a similar function. The RCS is a synthetic Enolase II RNA derived by IVT that, similarly to the IVT transcriptomes, is devoid of RNA modifications. Despite the reduced diversity of sequence in the RCS data, the overall results were comparable to those from the IVT transcriptome data (**Fig. S11A,B, Fig. 2A,B**), demonstrating that the RCS can be used as a “built-in” negative control for modification detection with DRS. Finally, we determined that basecall and modification probability distributions remain generally consistent between different Dorado models, although more m^6^A and Ψ sites are detected with high modification probability in Dorado model v5.1.0 compared to v5.0.0 (**Fig. S11C-E**).

Based on our observations, we propose a base probability cutoff of 0.8 and a modification probability cutoff of 0.98 for m^6^A (and 0.99 for Ψ and inosine) to reduce background signal.

### Filtering by predicted m^6^A/A proportion further reduces impact of background signal

We next examined the predicted m^6^A/A proportion distributions for all m^6^A sites in both modification probability-unfiltered and filtered NHDF datasets (**Fig. S12**). In addition to applying base and modification probability cutoffs, we restricted analyses to sites corresponding to adenosine residues in the reference sequence used for alignment, an approach also reported by Esfahani and colleagues (Ghohabi Esfahani et al. 2026) thereby further reducing the likelihood of background signal. We additionally required a minimum valid coverage of 20 reads per site, which is also consistent with the criteria applied in the same study. Valid coverage was defined as the sum of reads supporting the canonical adenosine base and reads supporting an m^6^A modification call (with ≥0.98 modification probability score). Unfiltered datasets showed that Dorado identifies 11.5x more m^6^A sites with the all-context model than with the DRACH model (**Supplementary Table 2**), consistent with the prior study by Cruciani and colleagues (Cruciani et.al. 2026), reporting 20x more all-context m^6^A sites than DRACH-context. Furthermore, we showed that the vast majority of m^6^A sites, 92% for all-context and 75% for DRACH-context, have predicted m^6^A/A proportion values <10%, with less than 1% of sites showing high (>50%) predicted proportions.

A similar analysis of the IVT transcriptome revealed large numbers of m^6^A sites derived from background signal with predicted m^6^A/A proportion values below 10%, most of which were eliminated by application of our filters (**Fig. S13A,B**). However, the remaining presence of low-proportion sites derived from background signal indicated that an additional layer of filtering to exclude sites with <10% proportion is beneficial. Notably, this filter was also adopted in the benchmarking study by Zou and colleagues (Zou et al. 2025). An additional argument supporting our filtering strategy is the observation that the effect of treatment with the METTL3-specific inhibitor STM2457 is much more pronounced at m^6^A sites with ≥10% predicted m^6^A/A proportion (**Fig. 2C,D; Supplementary Table 2**), regardless of the motif context. Applying modification probability and per-site proportion cutoffs substantially reduced the difference between the numbers of m^6^A sites detected with all-context and DRACH-context modes of Dorado (**Supplementary Table 2**). Application of our modification probability filtering strategy removed over 70% of the sites (**Fig. 2E**), regardless of the context, almost all of which showed predicted m^6^A/A proportions of <10%, without significantly changing the numbers of high-proportion sites detected. Similar results were also observed for the transcriptome-aligned NHDF and HD10.6 distributions (**Fig. S14, Fig. S15; Supplementary Table 2**).

To determine whether our filtering strategy could be applied to other RNA modifications detected by Dorado, we examined the distributions of predicted Ψ/U and inosine/A proportions in human genome-aligned DMSO-and STM2457-treated NHDF and HD10.6 datasets, before and after applying the ≥0.99 modification probability threshold. Similar to m^6^A (**Fig. S12**), the majority of detected Ψ and inosine sites exhibited low predicted modification proportions, with 92% of Ψ sites and 96% of inosine sites showing proportions below 10% (**Fig. S16 for NHDFs, Fig. S17 for HD10.6 cells**). This predominance of low-proportion Ψ sites is consistent with a recent study using 2-bromoacrylamide-assisted cyclization sequencing (BACS), which reported that 82% of Ψ modifications in human mRNAs occurred at proportions below 20% (Xu et al. 2024).

Importantly, analysis of the corresponding IVT datasets further supported the use of the ≥0.99 modification probability threshold for Ψ and inosine (**Fig. S13C,D**). Applying this threshold substantially reduced the number of sites detected in the IVT controls compared with the unfiltered data. For inosine, predictions derived from background signal were present in the unfiltered IVT data even at high (>50%) predicted modification proportions and were markedly reduced following modification probability filtering (**Fig. S13D**). These findings indicate that stringent modification probability filtering is necessary to reduce background predictions for Ψ and inosine before interpreting their estimated per-site proportions.

We have thus demonstrated that setting thresholds for basecall modification probabilities provides a robust method for identifying and filtering against the influence of background signal.

### Ψ and inosine sites exhibit the highest degree of modification-specific basecalling mismatches

An important consideration when interpreting the predicted modification proportions is that these estimates do not account for all reads overlapping a given position. Modification proportions are calculated from reads in which the corresponding reference base is basecalled as the expected canonical nucleotide and is subsequently assigned as either canonical or modified (above the selected modification probability threshold). Reads in which the position is basecalled as a different nucleotide, contains a deletion, is assigned another modification, or does not meet the selected modification probability threshold are therefore excluded from the proportion calculation. For the DRACH-context m^6^A model, modification calling additionally depends on preservation of the required sequence context, such that reads containing a non-matching base within the motif may also be excluded. Consequently, modification-associated effects on canonical basecalling could introduce bias into the reported modification proportions.

To assess the extent of basecalling mismatches, we examined alternative nucleotide calls at identified modification sites in the filtered DMSO-treated NHDF and HD10.6 datasets and compared, for each site, the percentage of reads containing a non-reference basecall with the predicted modification proportion (**Fig. S18**). For most sites, non-reference basecalls represented <25% of overlapping reads. Sites at which ≥25% of reads contained a non-reference basecall were comparatively uncommon and were predominantly associated with low predicted modification proportions (<10%). We examined the composition of alternative nucleotide calls at these sites across the different modification types. Notably, in both NHDF and HD10.6 cells, C accounted for the majority of alternative basecalls at Ψ sites, whereas G predominated at inosine sites (**Fig. S18**).

These observations indicate that basecalling errors can exclude a subset of reads from modification proportion calculations, with the extent and nature of this effect differing between modification types. The predicted modification proportions may therefore underestimate the true proportion of modified molecules at reported sites, as reads containing a modification but basecalled as an alternative nucleotide would not contribute to the modification estimate. This dependency may also result in some modification sites being missed altogether when basecalling errors reduce the number of reads eligible for downstream modification calling. Thus, the reported proportions should be interpreted as only estimates derived from reads satisfying the respective basecalling and modification-calling criteria.

### RNA modification detection rates differ between Dorado models

Dorado basecalling relies on models generated using Remora (github.com/nanoporetech/remora) that are frequently updated. This raises an important question as to whether these updates produce notably different outputs that could impact biological interpretation. To address this, we utilized multiple Dorado versions and Remora models to perform modification calling of our NHDF dataset. We restricted our analyses to m^6^A and Ψ as these are the only modifications that could be detected in previous models. Here, we observe that predicted m^6^A and Ψ proportion distributions are generally consistent between different Dorado versions and models (**Fig. S19**), albeit with greater numbers of low predicted proportion (< 10%) sites detected in older releases. There is a large overlap of the detected ≥ 10%-proportion of m^6^A (DRACH and all-context) and Ψ sites between versions/models, although there are a lot of unique all-context m^6^A and Ψ sites identified with just model v5.0.0 and not v5.1.0, that are reduced after applying our modification probability filter (**Fig. S20A,B,C**). Ψ sites uniquely detected with different models are found in slightly different sequence contexts (**Fig. S20C,D**). Thus, while RNA modification detection with Dorado is sensitive to changes in the underlying models/versions, this can be effectively controlled through the application of filters that only retain sites with high-confidence modification probability calls.

### GLORI and Dorado converge at sites with high predicted modification proportions

To assess the agreement of DRS Dorado-derived predictions with an orthogonal m^6^A detection method, we conducted GLORI on the same NHDF RNA preparations. To ensure a fair comparison of m^6^A sites across the two methods, we applied the following filtering strategies: For Dorado (DRACH model), this included thresholds on modification probability (A ≥ 0.8, m^6^A ≥ 0.98) and read coverage (≥ 20). For GLORI, we used filters based on coverage (≥ 20) and statistical significance (p-value ≤ 0.05). Next, we extracted m^6^A sites that were present in both DMSO and STM2457-treated samples and removed all sites with ≤ 10% predicted proportion in DMSO (**Fig. 3A**). This stepwise strategy reduced background signal and enabled a fair comparison of the high-confidence m^6^A sites identified by both approaches.

**Figure 3.**
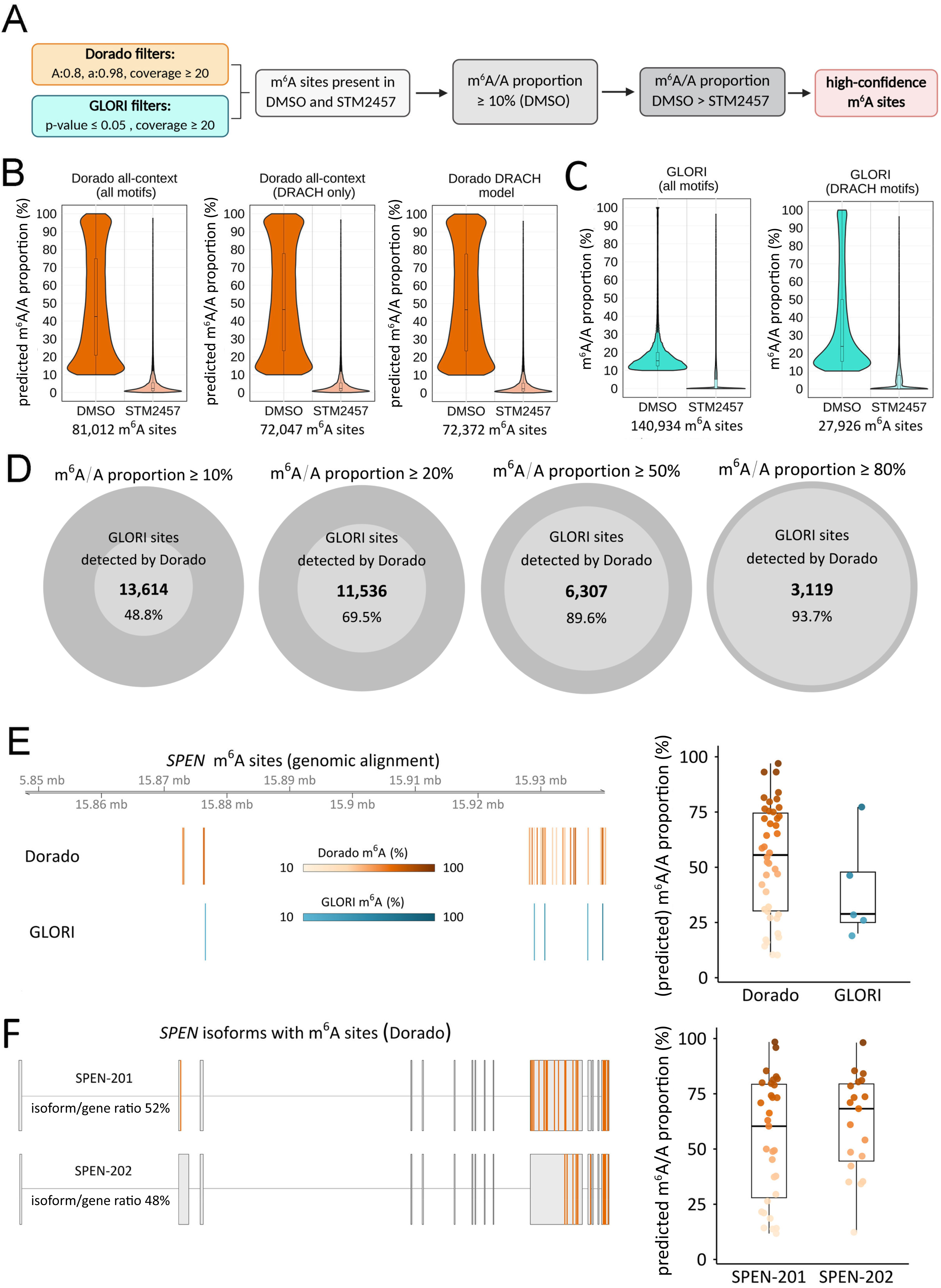
Comparison of m^6^A sites detected by Dorado and GLORI. **(A)** Filtering strategy for Dorado vs GLORI to obtain a high-confidence list of m^6^A sites in NHDFs. **(B)** Predicted m^6^A/A proportion distributions are plotted from genome-aligned DMSO-or STM2457-treated NHDF datasets using the indicated Dorado models and the filtering strategy shown in (A). **(C)** Predicted m^6^A/A proportion distributions from GLORI performed on DMSO-or STM2457-treated NHDFs are plotted, using the filtering strategy shown in (A). In addition, m^6^A sites were filtered for DRACH motifs (right plot). **(D)** Overlap of high confidence m^6^A sites detected by Dorado vs GLORI. GLORI and Dorado datasets were filtered using increasing predicted m^6^A/A proportion cutoffs (from left to right). GLORI sites overlapping with Dorado-identified sites are shown in light grey, whereas sites unique to GLORI are shown in darker grey. **(E)** m^6^A sites in the *SPEN* gene detected by GLORI and Dorado. Boxplots (right) represent the predicted m^6^A/A proportions per m^6^A site shown on the left. **(F)** *SPEN* isoforms with m^6^A sites, determined by Dorado. Boxplots (right) show the distribution of predicted m^6^A/A proportions on the m^6^A sites on *SPEN-201* and *SPEN-202*.

As shown in the violin plots, Dorado identified similar numbers of m^6^A sites using all-context and DRACH-context modification calling (81,012 and 72,372 sites, respectively), with predicted modification proportions strongly reduced upon STM2457 treatment in both cases (**Fig. 3B**). The two Dorado approaches differ in the sequence context in which m^6^A modifications are called: all-context calling permits m^6^A prediction at all adenosines, whereas DRACH-context calling restricts predictions to adenosines within DRACH motifs (DRACH, where D denotes A, G or U; R denotes A and G and H denotes A, C or U). Motif analysis of sites identified using all-context calling revealed nearly 9,000 sites outside DRACH motifs. Accordingly, when all-context calls were restricted to DRACH motifs, the resulting number of sites was similar to that obtained using DRACH-context calling (**Fig. 3B**). Unexpectedly, GLORI detects a substantially higher number of sites when using all motifs (140,934) compared to DRACH motifs alone (27,926) (**Fig. 3C**). The predominance of non-DRACH sites among the additional GLORI calls may reflect a higher contribution of background signal. Consistent with this interpretation, the overlap between Dorado and GLORI datasets remains virtually unchanged when comparing GLORI “all motif” and “DRACH motifs” (n=13,614) site sets, underscoring that the additional non-DRACH sites in GLORI (n= 113,008) contributed little to the observed overlap (**Fig. S21A**).

We obtained similar results when comparing m^6^A sites obtained with Dorado and GLORI from HD10.6 cells. Overall, we detected a similar number of m^6^A sites in HD10.6 cells using the all-context and DRACH-specific Dorado models (77,728 and 67,222, respectively) (**Fig. S21B**). By contrast, we detected 174,282 vs. 27,965 m^6^A sites using GLORI without and with DRACH filtering, respectively (**Fig. S21C**).

To evaluate how predicted m^6^A/A proportion thresholds affect the agreement between GLORI and Dorado, we compared the overlap of high-confidence GLORI-identified sites with Dorado m^6^A (DRACH context) calls across increasing proportion cutoffs (**Fig. 3D**). At the lowest threshold applied (≥ 10%), 13,614 sites overlapped, representing less than half (48.8%) of GLORI sites. However, as the stringency increased, the relative overlap increased sharply: at ≥ 20%, nearly 70% of GLORI sites overlapped (11,536 sites), at ≥ 50% almost 90% overlapped (6,307 sites), and at the highest cutoff (≥ 80%), over 93% of GLORI sites (3,119 sites) were also detected by Dorado. These results demonstrate that, while GLORI detects a broad set of potential m^6^A sites, Dorado increasingly recovers GLORI sites with higher m^6^A/A proportions.

GLORI relies on the high-efficiency conversion of adenosine to inosine, with overall gene A-to-I conversion rates needing to exceed 95-98% (Xu et al. 2024; Xie et al. 2025). Similarly, we observed the conversion rates for our NHDF and HD10.6 datasets to be 95% (**Supplementary Table 3**), but we found that small variations in gene conversion rates can strongly influence the sensitivity and specificity of GLORI, as only high-proportion m^6^A sites overlapped with DRS-derived predictions (**Fig. 3D**). When we analyzed the distribution of read coverage among m^6^A sites detected by both Dorado and GLORI or by GLORI alone (all motifs and DRACH motifs), we did not observe a striking difference. This suggests that read coverage does not account for the large number of sites detected only by GLORI (**Fig. S21D**). In contrast, comparison of m^6^A/A proportion distributions revealed an enrichment of high-proportion m^6^A sites among shared sites, whereas GLORI-only sites were predominantly in the 10-20% range (**Fig. S21E**). At low m^6^A/A proportions, the overall gene conversion rate may make it more difficult to distinguish background from true signal. In summary, given the technical limitations of the GLORI dataset, it provides complementary, rather than complete, orthogonal validation.

### DRS coupled with Dorado enables m⁶A prediction at RNA isoform level

Next, we wanted to highlight the advantage of long-read sequencing in terms of mapping reads with isoform-precision to the human transcriptome, which is not possible using GLORI due to its reliance on short-read sequencing. We first confirmed that mapping Dorado-basecalled sites to the human transcriptome results in the detection of similar numbers of m^6^A sites in both NHDFs and HD10.6 cells (**Fig. S21F,G**). To assess if DRS coupled with Dorado identifies distinct m^6^A profiles on different transcript isoforms, we analyzed the *SPEN* gene locus, which was previously shown to contain more than 100 m^6^A sites (Liu et al. 2023). On a genomic level, we detected 42 m^6^A sites using DRS, while GLORI only detected 5 m^6^A sites (**Fig. 3E**). Out of the 7 total isoforms of *SPEN*, *SPEN-201* and *SPEN-202* were detectable in DRS-derived data, considering a minimal coverage of 20 (**Supplementary Table 4**). We found that both isoforms were expressed at equal ratios in our NHDFs. When plotting the m^6^A sites detected by transcriptome-aligned DRS data, we found that the *SPEN-201* isoform exhibited more m^6^A sites, albeit with a lower median predicted proportion compared to the *SPEN-202* isoform (**Fig. 3F**). To further illustrate the ability of DRS to resolve isoform-specific m^6^A patterns, we included three more examples in the supplementary data. The monoexonic *KRTAP1-5*, with a single detectable isoform, demonstrates that DRS identifies more sites than GLORI, especially those with lower predicted m^6^A/A proportions (**Fig. S22A**). For *HSPA8*, which has 5 detectable m^6^A-containing isoforms, DRS reveals additional m^6^A sites present in two isoforms that cannot be resolved by GLORI (**Fig. S22B**). Similarly, *CITED2*, with three detectable m^6^A-containing isoforms, exhibits distinct isoform-specific m^6^A modification patterns detectable only by DRS (**Fig. S22C**). These examples highlight that DRS provides RNA-variant-resolved predictions that are not readily assignable using short-read GLORI data, as illustrated at several loci.

### Dorado profiling reveals m^6^A differences between fibroblast and neuronal datasets

To assess the reproducibility of Dorado across DRS datasets derived from different cell types, we compared the m^6^A methylomes of NHDFs and HD10.6 cells. To ensure a fair comparison, we extracted genes that were expressed in both cell types. Next, we compared the total numbers of high confidence m^6^A sites in NHDFs and HD10.6 cells within the genes expressed in both cell types and found that 87% of HD10.6-derived m^6^A sites overlap with those detected in NHDFs (**Fig. 4A**). Site-specific predicted m⁶A/A proportions from the shared gene set correlated strongly across cell types (Pearson r = 0.93), although we observed a global shift in the HD10.6 dataset, where predicted m⁶A/A proportions were generally higher than in NHDFs (**Fig. 4B**). Plotting the m^6^A/A proportions of the 38,217 m^6^A sites shared between both cell types showed a strong increase in the median predicted m⁶A/A proportions of HD10.6 cells (70.2%), compared to NHDFs (54%) (**Fig. 4C**).

**Figure 4.**
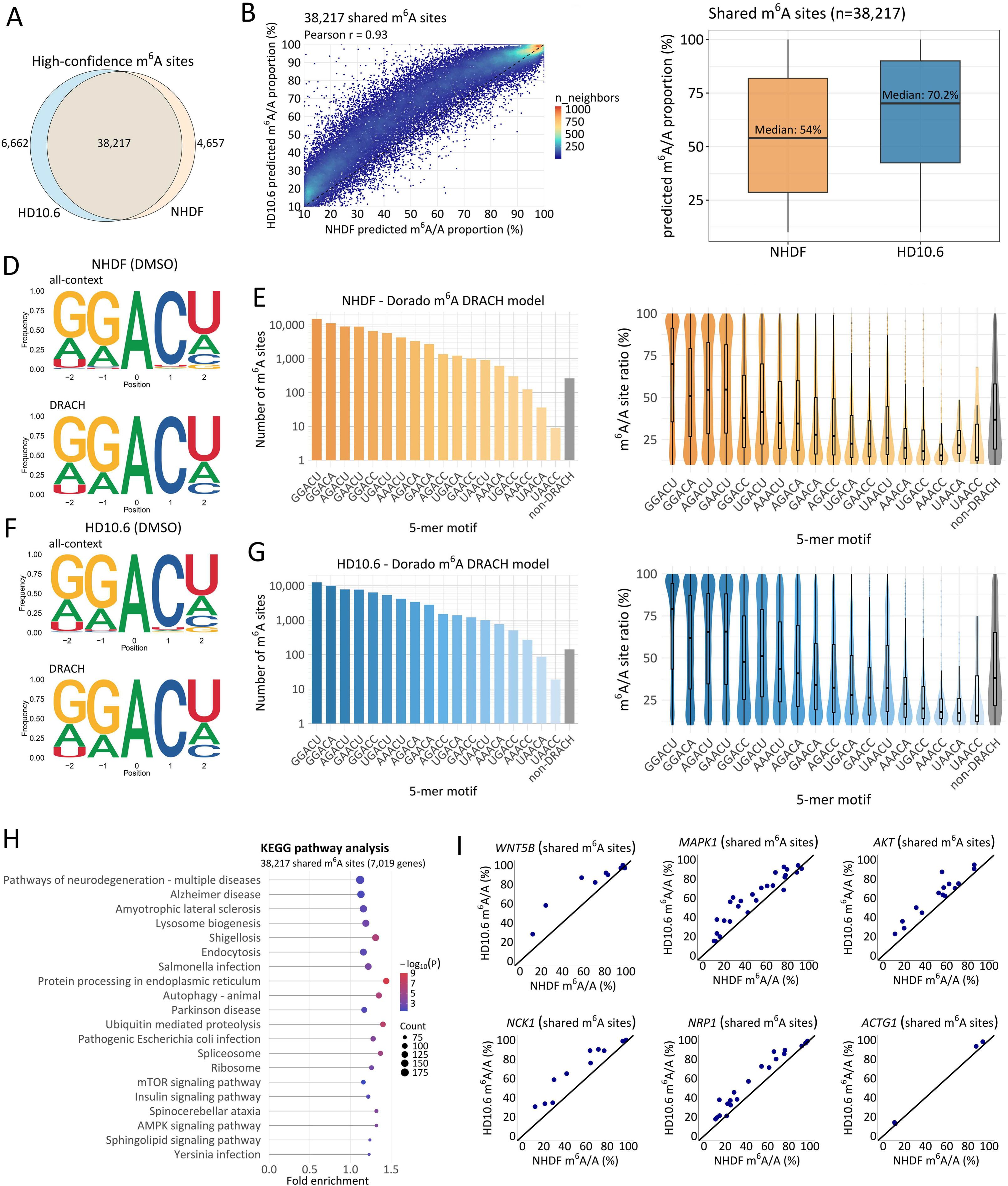
Comparative analysis of m⁶A modification patterns in NHDFs and HD10.6 cells using Dorado v0.9.0. **(A)** Overlap of high-confidence m⁶A sites between NHDFs and HD10.6 cells. **(B)** Correlation of predicted m^6^A/A proportions at shared m^6^A sites (n=38,217). **(C)** Boxplots showing the predicted m^6^A/A proportion distributions of the 38,217 detected in both NHDFs and HD10.6 cells. **(D,E)** Motif analysis showing m^6^A (all-context and DRACH-context) motif logos, 5-mer distributions and predicted modification proportions in NHDFs. **(F,G)** Same as in (D,E) but for HD10.6 cells. **(H)** KEGG pathway analysis (DAVID Bioinformatics Resources) of shared m^6^A-containing genes. **(I)** Predicted m^6^A/A proportions of representative examples of individual genes from enriched pathways (H). *ACTG1* serves as a housekeeping control.

We next aimed to conduct a comparative motif analysis of m⁶A sites detected in NHDFs and HD10.6 cells. Sequence logos confirmed strong enrichment of the canonical DRACH consensus motif for both cell types in modification probability-filtered data, with the GGACU pentamer being the most prominent (**Fig. 4D,E**). Quantification of 5-mer distributions revealed highly similar motif preferences across datasets. While the global motif architecture of m⁶A deposition is conserved between NHDFs and HD10.6 cells, we observed a trend toward higher predicted m^6^A/A proportions in the HD10.6 dataset, particularly within the most abundant DRACH motifs (**Fig. 4F,G**).

Intriguingly, even after filtering, Dorado called m^6^A at more than 100 motifs that deviate from the canonical DRACH consensus. To understand what these represent, we examined the 15 most frequently detected non-DRACH motifs in NHDFs using the DRACH model. AAAAC and AAAAA were the most abundant, followed by ACACU and GGACG. Most sites within these motifs showed a strong reduction in predicted m^6^A/A proportions upon STM2457 treatment, indicating that they were deposited by METTL3 (**Fig. S23A**). Within the top 15 motifs, we found two motifs that conformed to DRAC but not DRACH. Because DRAC sites have been reported as authentic m^6^A sites (Körtel et al. 2021; Zhou et al. 2024; Kang et al. 2025), these calls are likely biologically real (**Fig. S23A**). For the remaining motifs, the discrepancy that they were sensitive to STM2457 treatment but neither in a DRACH or DRAC context prompted us to ask whether these calls instead reflect systematic limitations of nanopore sequencing and basecalling. Many non-DRACH sites lie one or two nucleotides from a genuine DRACH motif. For example, m^6^A sites within ACACU motifs are located in proximity to DRACH sites, shifted by one or two positions (**Fig. S23B**). Another category that contributes to the detection of non-DRACH sites are single nucleotide polymorphisms (SNPs). One example is represented by the GGACG motif, in which the final G position of the 5-mer is basecalled as A in 49% and G in 51% of all reads. This makes the sequence context a DRACH motif in half of the reads, and a non-DRACH motif in half of the other reads (**Fig. S23C**). Another category reflected in the non-DRACH motifs are sites that lie within homopolymer sequences. For example, several sites within AAAAA and AAAAC are reported by Dorado, but because the nanopore signal is poorly resolved within homopolymers, Dorado appears unable to localize the modified adenosine precisely, assigning it to neighboring positions rather than to the adjacent DRACH motif (**Fig. S23D**).

In parallel, we investigated the occurrence of non-DRACH sites using the all-context model, and as expected, Dorado predicted more than 3,000 sites. Among the top 15 motifs, GGACG, GUACU, AGACG and GGAUU were most abundant (**Fig. S24A**). As with the DRACH model, GGACG calls derived largely from SNPs or represented genuine m^6^A sites in a DRAC context. GGAUU, the second most abundant motif, behaved differently, as we found no evidence of miscalling or problematic sequence context. m^6^A sites within this motif were STM2457-sensitive, suggesting authentic METTL3-dependent methylation, however, this requires further validation (**Fig. S24B**). Notably, the all-context calls separated into two classes: i) STM2457-sensitive sites and ii) STM2457-insensitive sites (**Fig. S24A**). The insensitive sites were enriched for motifs resembling the METTL16 consensus. ACAGA is a clear example, recurring in *MAT2A*, which is a well-established METTL16 target. Their insensitivity to METTL3 inhibition is consistent with METTL16-dependent deposition, though direct validation, for example in METTL16-depleted cells, will be required to confirm this (**Fig. S24C**).

Metagene analysis showed, as known previously, that m^6^A is enriched near the stop codons and in the 3’ UTRs (**Fig. S25A**). Several studies have examined the relationship between exon architecture and m^6^A distribution on mRNAs (Yang et al. 2022; He et al. 2023; Luo et al. 2023), and so we also examined m^6^A sites in monoexonic transcripts separately and found that monoexonic transcripts are more broadly m^6^A-modified across the entire CDS (**Fig. S25A**). m^6^A (both all-context and DRACH-context) distribution profiles look less typical for sites with predicted m^6^A/A proportions lower than 10% in both NHDF and HD10.6 datasets (**Fig. S25B,C**). For all-context m^6^A (**Fig. S25B**), the modification probability filter also proves to be deciding for the distribution profile, together with predicted m^6^A/A proportion of ≥ 10%.

To connect shared m⁶A sites with their functional context and illustrate their distribution on individual transcripts, we conducted a KEGG pathway analysis with the genes that contain shared m^6^A sites between NHDFs and HD10.6 cells. We used an overlap of all genes that were expressed in both cell types as a background gene set. We observed a significant enrichment in pathways related to neurodegeneration, infection, and intracellular signaling (**Fig. 4H**). To further illustrate these findings, we examined representative genes contained in one or more of these pathways. *WNTB5*, *MAPK1*, *AKT1*, *NCK1,* and *NRP1* all showed robust methylation in both NHDFs and HD10.6 cells, with consistently higher predicted m^6^A/A proportions observed in the neuronal-like context. In contrast*, ACTG1*, included as a housekeeping control not represented in the enriched pathways, displayed stable and comparatively uniform methylation across both cell types (**Fig. 4I**).

Together, these results demonstrate that while the global architecture of m^6^A deposition is largely conserved between primary fibroblasts and neuronal-like cells, neuronal-like cells exhibit globally higher predicted m^6^A/A proportions, a difference that is readily detectable in individual genes. However, the biological significance of these differences remains unclear, and whether they are associated with differences in RNA abundance, stability, translation, or other cell type-specific features will require further functional investigation.

## Discussion

High rates of false positives remain a major problem in epitranscriptomic mapping tools, even with the advent of modification-aware basecalling models (Cruciani et al. 2025). Comparing the outputs of orthogonal methods performed on the same input material, combined with the use of IVT RNA and/or specific methyltransferase inhibition, offers an effective mitigation strategy (Cruciani et al. 2025; Diensthuber and Novoa 2025). The results presented here establish a robust framework that minimizes background signal and enables isoform-specific and quantitative m^6^A mapping. Through a direct comparison with GLORI, we showed that combining DRS datasets with Dorado modification-aware basecalling provides greater scalability, isoform resolution, and reproducibility across different cell types. Further, the simplicity of DRS library preparation enhances consistency, whereas GLORI is constrained by a laborious and technically challenging protocol that is extremely sensitive to adenosine conversion rates (Shen et al. 2024). Despite these limitations, concordance with Dorado increased at higher GLORI-reported m^6^A/A proportions, with the majority of high-proportion GLORI sites also detected in our filtered DRS datasets. By expanding our analysis across multiple human cell types, we further determine that the sites of m^6^A installation are conserved for the most part but that the predicted m^6^A/A proportions show cell-type-specific differences.

Crucially, the inclusion of transcriptome-wide IVT datasets and the use of a selective METTL3 inhibitor (STM2457) provide a robust approach to developing filters that dramatically decrease m^6^A detection arising from background signal. The impact of both approaches was readily visualized by plotting modification probability distributions, enabling our determination that retaining only m^6^A calls with confidence scores of 0.98 or greater retained almost all STM2457-sensitive m^6^A sites while eliminating large numbers of m^6^A sites derived from background signal reported for the IVT datasets. This approach is supported by a recent study that highlighted that Dorado introduces large numbers of false positives outside canonical DRACH contexts (Xie et al. 2025). Our data confirm that in the unfiltered datasets, the number of m^6^A sites derived from background signal is higher in the all-context model compared to the DRACH model. This may reflect the degenerate nature of the canonical DRACH consensus and understudied differences in sequence preference between different cell types and organisms. Retaining only sites at which predicted m^6^A/A proportions exceeded 10% further reduced the impact of background signal while also focusing the results on identification of sites that are more likely to be biologically significant. The combined effect of these filters was to provide a strong agreement between the m^6^A all-context and m^6^A DRACH model in the detection of high-confidence m^6^A sites, as observed by filtering for DRACH motifs from m^6^A sites detected with the all-context model (**Fig.3B**).

Prior studies have emphasized the need for isoform-resolved m^6^A quantification by demonstrating widespread heterogeneity in m^6^A levels between genes and their isoforms (Guo et al. 2025; Xie et al. 2025). Here, we showed that aligning DRS datasets to the human transcriptome reveals differential m^6^A installation patterns between isoforms, underscoring the importance of accurate, isoform-specific m^6^A quantification at single-molecule resolution and highlighting the advantage of DRS against short-read sequencing approaches such as GLORI (**Fig. 3E,F**, **Fig. S22**).

Recently, the ability of Dorado and m6Anet (Hendra et al. 2022) to identify m^6^A in DRS datasets from 293T and HeLa cells was systematically evaluated using IVT RNA and comparisons to published datasets obtained by GLORI and eTAM-seq (Zou et al. 2025). In this case, the authors found that Dorado (v0.7.2) outperforms m6Anet but also advocated for the importance of applying filters to reduce false-positive calls. Likewise, we found that IVT RNA served as a valuable negative control to detect background signal. Furthermore, using the RNA calibration strand (RCS) as a negative control provided a convenient and simpler alternative to IVT RNA, yielding comparable probability distributions and thereby enabling a built-in control for routine experiments (Viehweger et al. 2019). While m^6^A detection is generally reproducible across different Dorado modification-aware basecalling models, model-specific differences are present, i.e., model v5.0.0 produces substantially more low-confidence all-context m⁶A and Ψ calls than v5.1.0 (**Fig. S11C-E**), highlighting that the models remain under active development and that benchmarking should be repeated when switching to a new model. Recent work further highlights that continued improvements in both nanopore sequencing chemistry and basecalling models significantly enhance RNA modification detection accuracy, emphasizing the importance of revisiting benchmarking and filtering strategies as the models evolve (Hewel et al. 2025).

Notably, Dorado v1.1.0 was included in a systematic comparison of computational approaches for the detection of six RNA modification types (Luo et al. 2026). While this study provides an important broad assessment of currently available computational methods, the workflow and filtering criteria used to derive high-confidence modification sites from Dorado predictions were not explicitly described. Our findings demonstrate that appropriate filtering and the inclusion of modification-free controls substantially reduce Dorado predictions derived from background signal. Thus, broader comparisons of modification-detection tools would benefit from consideration of the criteria used to define modification sites, particularly as Dorado models continue to evolve. In this respect, the framework established here complements such benchmarking efforts by providing practical guidance for obtaining high-confidence modification calls from Dorado output.

Focusing on other modifications that are detected by Dorado, we observe that stringent probability and predicted m^6^A/A proportion filtering substantially reduces background predictions for inosine and Ψ, with similar filtering strategies also supported by recent studies (Verstraten et al. 2025; Ghohabi Esfahani et al. 2026). For inosine, we observed an enrichment of high-probability (> 0.99) inosine sites in genome-aligned data from STM2457-treated samples. Further studies are needed to determine whether this reflects increased intron retention or misclassification of modifications by the current inosine model. Nevertheless, very recent systematic benchmarking of Dorado modification-aware basecalling models highlights important limitations that warrant caution when interpreting these predictions (Diensthuber et al. 2026). Modification-aware models in particular, were shown to exhibit cross-reactivity with chemically related modifications, including substantial cross-reactivity of the Ψ model with m^1^Ψ and of the m5C model with hm^5^C and ac^4^C. Moreover, neighboring modifications can alter the nanopore signal and contribute to predictions arising from background signal. Thus, even high-confidence predictions cannot necessarily be considered unambiguous evidence of the corresponding chemical modification, particularly in the absence of orthogonal validation. These findings further emphasize the importance of modification-free controls and stringent filtering, while also highlighting the need for continued improvement and benchmarking of modification-aware basecalling models.

Emerging evidence indicates that m⁶A can occur outside the DRACH consensus *in vivo*, and our attribution of most non-DRACH GLORI calls to background signal should not be interpreted as a claim that cellular m⁶A is restricted to this motif. Our dissection of non-DRACH DRS calls in the filtered datasets revealed that technical and biological explanations coexist. Several classes were clearly artefactual, arising from positional imprecision relative to neighboring DRACH sites, from SNPs that render the context DRACH in only a fraction of reads, or from homopolymeric stretches in which the modified adenosine cannot be localized unambiguously (**Fig. S23, Fig. S24**). However, we also found two abundant motifs that conformed to DRAC, but not DRACH, consistent with reports of authentic m⁶A at DRAC sites, and STM2457-sensitive GGAUU sites showed no evidence of miscalling, which reflects previous findings (Körtel et al. 2021; Zhou et al. 2024; Kang et al. 2025). A recent study showing measurable activity of recombinant METTL3-METTL14 on non-DRACH substrates in vitro further supports our interpretation that the observed GGAUU sites represent valid non-canonical m^6^A sites (Zhang et al. 2024). Conversely, a distinct class of STM2457-insensitive sites was enriched for METTL16-like motifs, including recurrent ACAGA sites in *MAT2A* (Su et al. 2022), pointing to writers other than METTL3 rather than to error. Our results suggest that DRS is in principle well suited to detecting non-consensus m⁶A sites, as we included STM2457, which readily makes METTL3-dependent vs. -independent sites discernible. Establishing individual non-canonical sites nonetheless requires read-level inspection of the surrounding sequence, awareness of local genetic variation, and future orthogonal validation.

While DRS is the only approach that allows for robust isoform-specific m^6^A mapping, comparisons with orthogonal approaches remain essential to validate findings. When comparing Dorado and GLORI using the same input RNA, we unexpectedly observed that only 50% of m^6^A sites with predicted proportions of above 10% that were identified by Dorado were also present in the GLORI data, while Dorado predictions captured an increasing proportion of GLORI sites detected at high m^6^A/A proportions (**Fig.3D**). One factor that may contribute to this discrepancy is the sub-optimal gene-wide adenosine conversion rate of approximately 95% achieved in our GLORI datasets, compared with the 98–99% reported in the original study (Shen et al. 2024). Incomplete adenosine conversion could increase background signal and potentially reduce the ability to distinguish m^6^A sites with lower predicted proportions. The large number of GLORI-detected sites outside DRACH motifs may similarly reflect increased background signal (**Fig. 3C**), although differences in detection sensitivity, sequencing depth, and other methodological biases cannot be excluded based on the current analyses. Further methodological updates to GLORI (Sun et al. 2025) may help improve its feasibility and facilitate more robust comparisons with DRS.

Prior analyses of predicted m^6^A/A proportion distributions across immortalized human cell lines, including cancer cell lines, have revealed broadly similar m^6^A proportion distributions (Guo et al. 2025). By contrast, our analysis of primary fibroblasts (NHDFs) and neuronal cells (differentiated HD10.6 cells) revealed an overall increase in predicted m^6^A/A proportions in HD10.6 cells compared to NHDFs, especially in genes associated with neuronal function. While further studies are warranted to confirm and investigate the underlying mechanisms and functional consequences of these differences, this aligns with the importance of m^6^A in neuronal function (Yu et al. 2021). This includes m^6^A maps from synaptic and nuclear RNA isolated from the mouse dorsal hippocampus, which showed that synapse-specific RNAs had generally higher methylation levels (Sun et al. 2025). Intriguingly, neuronal regulation pathways utilize differentially spliced isoforms (Su et al. 2018), and a recent study revealed isoform-specific differences in m^6^A sites and predicted m^6^A/A proportions in the human brain that are linked to m^6^A machinery expression and cell-type context (Gleeson et al. 2025).

In summary, using matched RNA samples from different experimental conditions and from different human cell types analyzed with both Dorado and GLORI orthogonal methodologies, we establish a robust pipeline for precise and quantitative transcriptome-wide detection of m^6^A in DRS datasets. Importantly, we recommend the inclusion of a modification-free control as a best practice for accurate and reliable DRS-based RNA modification detection. Future studies should aim at extending this approach using orthogonal sequencing approaches to benchmark Dorado-based detection of Ψ, m^5^C and inosine, as well as the newly released models for 2’-O-methylation.

## Methods

### Cell culturing and STM2457 treatment

NHDFs (Lonza) were cultured in Dulbecco’s Modified Eagle Medium (DMEM, Gibco), supplemented with 5% heat-inactivated FBS (HI-FBS, Gibco) and 1% penicillin-streptomycin (PS, Lonza) under standard culture conditions. The HD10.6 cell line was originally derived from fetal human dorsal root ganglia (DRG) and immortalized by overexpression of the v-myc oncogene under tetracycline regulation. Upon addition of tetracycline and neurotrophins, these cells can be differentiated into neuron-like cells that show typical neuronal morphology and nociceptive properties (Raymon et al. 1999). For our experiments, we used HD10.6 cells that have been matured for 10 days. For maintenance of undifferentiated HD10.6 cells (a gift from Anna Cliffe, University of Virginia School of Medicine), T-75 flasks were coated with 17 µg/mL fibronectin in PBS for 15 min at 37°C before adding the cell suspension. Cells were grown in Advanced DMEM/F12 (Gibco), supplemented with 1x Neurocult SM1 (STEMCELL Technologies), 2 mM L-glutamine (Gibco), 1x Primocin (Invivogen) and 10 ng/mL prostaglandin E1 (Sigma). 0.5 ng/mL bFGF (PeproTech) were added freshly to each media change. For maturation, HD10.6 cells were seeded at a density of 25.000 cells/cm² in 6-well plates which were pre-coated overnight with 50 µg/mL poly-L-ornithine hydrobromide (Sigma) in 0.5M borate buffer pH 8.5 (Boston Bioproducts), and 1 µg/mL fibronectin (overnight). After cell attachment overnight, media was changed to Neurobasal media (Gibco), supplemented with 1x Neurocult SM1, 2mM L-glutamine, 1x Primocin, 1 µg/mL doxycycline, 50 ng/mL NGF (Alomone Labs), and 25 ng/mL CNTF, GDNF and NT-3 (PeproTech). Media was changed every three days until HD10.6 cells showed complete neuronal morphology after 12 days. NHDFs and matured HD10.6 cells were treated with 30 µM STM2457 (Sigma) or DMSO for 48h, harvested in TRIzol (Invitrogen), and stored at -80°C until RNA isolation.

### Cell viability testing

Cell viability was assessed using the CellTiter-Glo assay (Promega), performed according to the manufacturer’s protocol. NHDF and HD10.6 cells were seeded in 96-well plates and treated with either DMSO or 30 µM STM2457 for 48h, using 10 wells per condition. Following treatment, the culture medium was aspirated and replaced with 50 µL of assay reagent. After a 2 min mixing step and a 10 min incubation at room temperature, luminescence was recorded using a PerkinElmer EnVision 2103 Multilabel Reader. Statistical significance was assessed using an unpaired, two-tailed Student’s t-test.

### RNA isolation

RNA was extracted by the TRIzol method according to the manufacturer’s instructions. Briefly, 0.2 mL chloroform per 1 mL TRIzol were added and mixed by vortexing. After centrifugation for 30 min at 12,000xg at 4°C, the aqueous phase was transferred to isopropanol (0.5 mL per 1 mL TRIzol), mixed and incubated for 10 min at room temperature. Subsequently, the samples were centrifuged for 10 min at 12,000xg at 4°C and the RNA pellets were washed twice in 75% fresh ethanol. After air-drying for 5-10 min, RNA was dissolved in nuclease-free H2O and stored at -80°C until further processing. One third of each sample was used for DRS and the remaining same-input material was processed for GLORI. For both DRS and GLORI, poly(A) RNA was selected using the Dynabeads™ mRNA Purification Kit (Invitrogen), according to the manufacturer’s instructions.

### GLORI-Sequencing

GLORI reactions were largely performed following the published protocol for RNA protection, deamination and deprotection with several minor adjustments (Shen et al. 2024). Briefly, poly(A) RNA was fragmented at 94°C for 2 min (NEB E6150S) and subjected to glyoxal protection, sodium-nitrite-mediated deamination (750 mM NaNO_2_ at 16°C for 8h and 4°C for 6h), and deprotection in triethylammonium acetate-formamide buffer at 95°C for 10 min. Library construction was conducted using a ligation-based strategy as described by Liu and colleagues incorporating 5’adapters with 11-nt UMIs (Liu et al. 2025).

Indexed libraries were amplified with NEBNext Dual Primers Set 2 (E7780S), purified using AMPure XP beads, and validated on an Agilent TapeStation (D1000 High Sensitivity). Sequencing was performed on an Illumina NovaSeq 6000 platform (150 bp paired-end mode; > 80 million reads per sample).

For data processing, we employed the same computational pipeline described in Liu et al (Liu et al. 2025), but implemented within a Snakemake-based framework that automates the complete analysis starting from raw fastq files (Köster and Rahmann 2012). Details on software versions, workflow availability, and repository links are provided in the Data and code availability section below.

### Generation of a transcriptome-wide IVT RNA library

We sequenced the IVT RNA from both NHDFs and HD10.6 cells along with the native poly(A) RNA to evaluate potential background signal influenced predictions by Dorado. This was achieved by converting the poly(A) RNA to cDNA using a poly(A) primer and a T7-incorporating strand switch primer and using the cDNA as a template for IVT synthesis.

### Nanopore Sequencing of poly(A) RNA

250-300 ng of isolated poly(A) RNA and 25 ng RNA calibration strand (RCS) were used as input for the standard DRS SQK-RNA004 protocol. Resulting libraries were loaded onto individual PromethION RNA flow cells before sequencing on the PromethION 2 Solo instrument for 48h.

### Nanopore basecalling

Basecalling was performed using Dorado versions v0.5.0, v0.6.0, v0.7.0, v0.8.0 and v0.9.0 (ONT) in super accuracy mode with modification detection enabled for m^6^A (DRACH context) in v.0.5.0 and 0.6.0 (model rna004_130bps_sup@v3.0.1_m6A_DRACH@v1) and 0.8.0 and 0.9.0 (model rna004_130bps_sup@v5.1.0), m^6^A (all context) in v0.7.0 (model rna004_130bps_sup@v5.0.0_m6A@v1),

0.8.0 and 0.9.0 (model rna004_130bps_sup@v5.1.0), Ψ in v0.7.0 (model rna004_130bps_sup@v5.0.0_pseU@v1), v0.8.0 and v0.9.0 (model rna004_130bps_sup@v5.1.0), and inosine and m^5^C in v.0.9.0 (model rna004_130bps_sup@v5.1.0). Each dataset was processed with adapter trimming. Resulting unaligned BAM files were converted to FASTQ files containing modification tags for every read in the FASTQ read headers, using SAMtools v1.18 (Li et al. 2009) [*samtools fastq -TMM,ML*].

### Genome and transcriptome level alignments

For genome-level alignment, reads were aligned against a combined human genome reference (GRCh38.p14), and the RCS gene sequence, using minimap2 v2.26 [*minimap2 -ax splice -k14 -y -uf --secondary=no*]. For transcriptome-level alignment, the reads were aligned against the human transcriptome (v47), obtained via GENCODE (Mudge et al. 2025) [*minimap2 -t 8 -ax map-ont -y -L -p 0.99*]. Using SAMtools v1.18, resulting genome-and transcriptome-aligned SAM files were converted to BAM files retaining only primary alignments [*samtools view -b -F2308* and *samtools view -b -F2324,* respectively]. BAM files were sorted [*samtools sort*] and indexed [*samtools index*].

### Subsampling aligned BAM files

Aligned BAM files were randomly subsampled to 200,000 reads using *reformat.sh* from BBMap v39.01 [samplereadstarget=200000]. Ten independent subsampling iterations were performed for each dataset using different random seeds (sampleseed=1-10).

### Processing of RNA modification calls using Modkit

Probability distributions of the detected sites in human genome-and transcriptome-aligned reads were generated using sample-probs subcommand from Modkit v0.4.1 [sample-probs --hist --only-mapped --percentiles 0.1,0.2,0.3,0.4,0.5,0.6,0.7,0.8,0.85,0.9 -f 0.1]. For RCS reads -f value was increased to 0.25. The resulting .tsv files were imported into RStudio (v. 2024.12.1+563) and plotted using a custom R(v. 4.4.2) script (Dorado_mod_probability_distributions.R), with the following packages: dplyr (v1.1.4), ggplot2 (v3.5.1), reshape2 (v1.4.4), patchwork (v1.3.0).

To generate extended bedMethyl files with m^6^A-modified and unmodified base counts from the reads mapping against the potential modification sites, we used Modkit pileup subcommand with the thresholds [*--filter-threshold A:0.8 --filter-threshold T:0.8 --filter-threshold C:0.8 --filter-threshold G:0.8 --motif A 0 --ref $REF*] with and without modification threshold filter [e.g. *--mod-thresholds a:0.98*] to generate our “filtered” and “unfiltered” datasets, respectively. Canonical nucleotide calls (A, C, G, U) were retained with > 80% probability scores for all modification datasets. Modification thresholds were defined by our statistical framework (see below) and were set at 98% for m^6^A and 99% for all the other modifications. *--motif* was adjusted to the base for which modification status was examined, which further restricted the analysis to only the positions that matched canonical nucleotides in the reference genome. The resulting sites were further filtered according to the read number per site (≥ 20 reads) for all the downstream analysis.

### Statistical framework for modification probability cutoff determination

Modification probability distributions were analysed using a two-stage statistical framework implemented in R to identify the probability threshold above which sites are considered genuinely modified relative to an unmodified in vitro transcribed (IVT) reference. For each sample, the per-bin log2 ratio of modification probability fractions was computed as log2(sample/IVT), with a pseudocount of 1×10⁻⁶ added to each value prior to transformation to avoid undefined values. This ratio quantifies fold-enrichment of the sample relative to IVT at each probability bin and is insensitive to differences in absolute signal magnitude across samples. For the first stage, change-point detection was applied to the log2 ratio series using the Pruned Exact Linear Time algorithm (Killick et al. 2012) implemented in *changepoint* R package (Killick and Eckley 2014), with a manual penalty value of 1.0 and a minimum segment length of 3 bins. The first detected change-point defines the end of the baseline region (i.e. the range of probability bins over which sample and IVT distributions are statistically indistinguishable) and the start of the signal region where enrichment above baseline begins. For the second stage, the modification probability cutoff was defined as the first bin at or after the first change-point at which the log2(sample/IVT) ratio exceeded a pre-specified minimum fold-enrichment threshold. This cutoff represents the lowest modification probability at which a site shows at least 1.5-fold more signal (i.e. log2 ratio > 0.585) than would be expected from an unmodified transcript. To characterise uncertainty in the log2 ratio estimates, bootstrap confidence intervals (95%, 2000 iterations) were computed by resampling bins with replacement and recalculating the log2 ratio distribution.

### KEGG pathway enrichment analysis

KEGG pathway over-representation analysis was performed using DAVID (Huang da et al. 2009). Genes harboring shared m⁶A sites in both NHDF and HD10.6 cells were used as input. All genes detected in both datasets were used as a custom background gene set (∼13,000 genes). Pathways were considered significant at a P-value ≤ 0.05. For visualization, the top 20 KEGG pathways containing the highest number of genes were selected.

### Analysis and visualization of predicted modification proportions and distributions

Histograms of predicted modification proportion distributions, as well as metaplots, were generated using RStudio and R v4.4.2 with the following libraries: data.table, patchwork, scales, tidyr, dplyr, ggplot2^4^. Access to scripts used for analysis and visualization is detailed below.

### Data and code availability

Raw fastq and pod5 datasets generated in this study are available via the European Nucleotide Archive (ENA) under the accession number PRJEB101380. R scripts used in the analysis and visualization of data presented herein are available in the https://github.com/DepledgeLab/HOLDEN repository. The Snakemake workflow implementing the GLORI analysis pipeline is available in the https://github.com/gp-micro/glorirepository.

## Acknowledgements and funding

We thank our colleagues in New York and Hannover for their continuous input on these studies and to Dr. Shino Murakami for access to an unpublished assay. DPD is supported by a German Centre for Infection Research (DZIF) Associate Professorship and the NIAID grants R01-AI170583 and R01-AI152543. DPD also receives funding from the Deutsche Forschungsgemeinschaft (DFG, German Research Foundation) under Germany’s Excellence Strategy -EXC 2155, project number 390874280. EL is supported by the NIAID grant R01-AI170583 as well as the Hannover Biomedical Research School (HBRS) and the Center for Infection Biology (ZIB). DO is supported by the Walter Benjamin Programme of the DFG, project 554758329. ACW is supported by NIAID grants R01AI176335 and R01-AI170583. https://github.com/nanoporetech/modkit.

---

**Figure S1.**
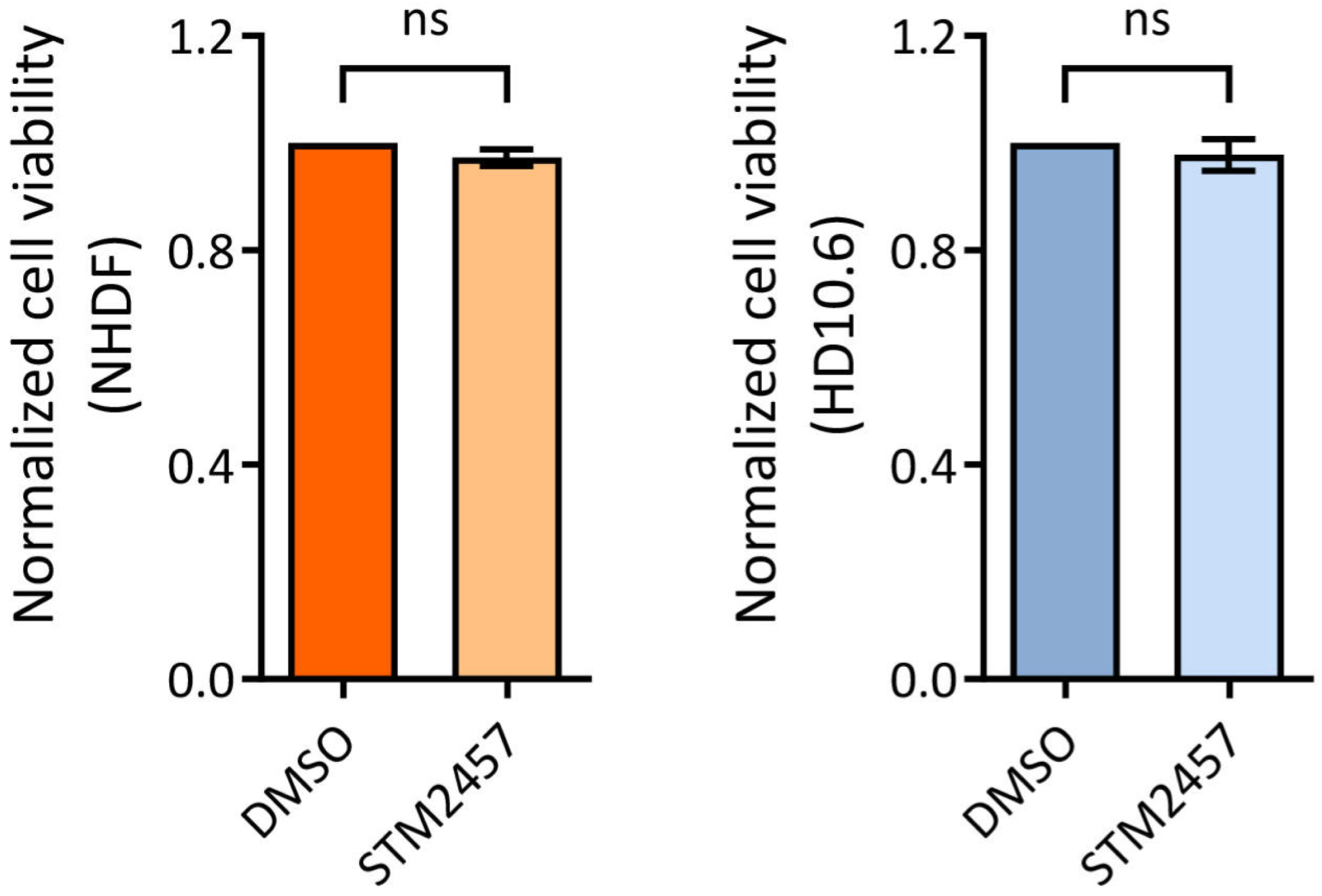
Cell viability assays in STM2457-treated NHDFs and HD10.6 cells. Cell viability was assessed in NHDF and HD10.6 cells after 48h of DMSO or 30 µM STM2457 treatment using CellTiter-Glo (Promega). Statistical significance was determined from 10 replicate wells per condition using an unpaired, two-tailed Student’s *t*-test.

**Figure S2.**
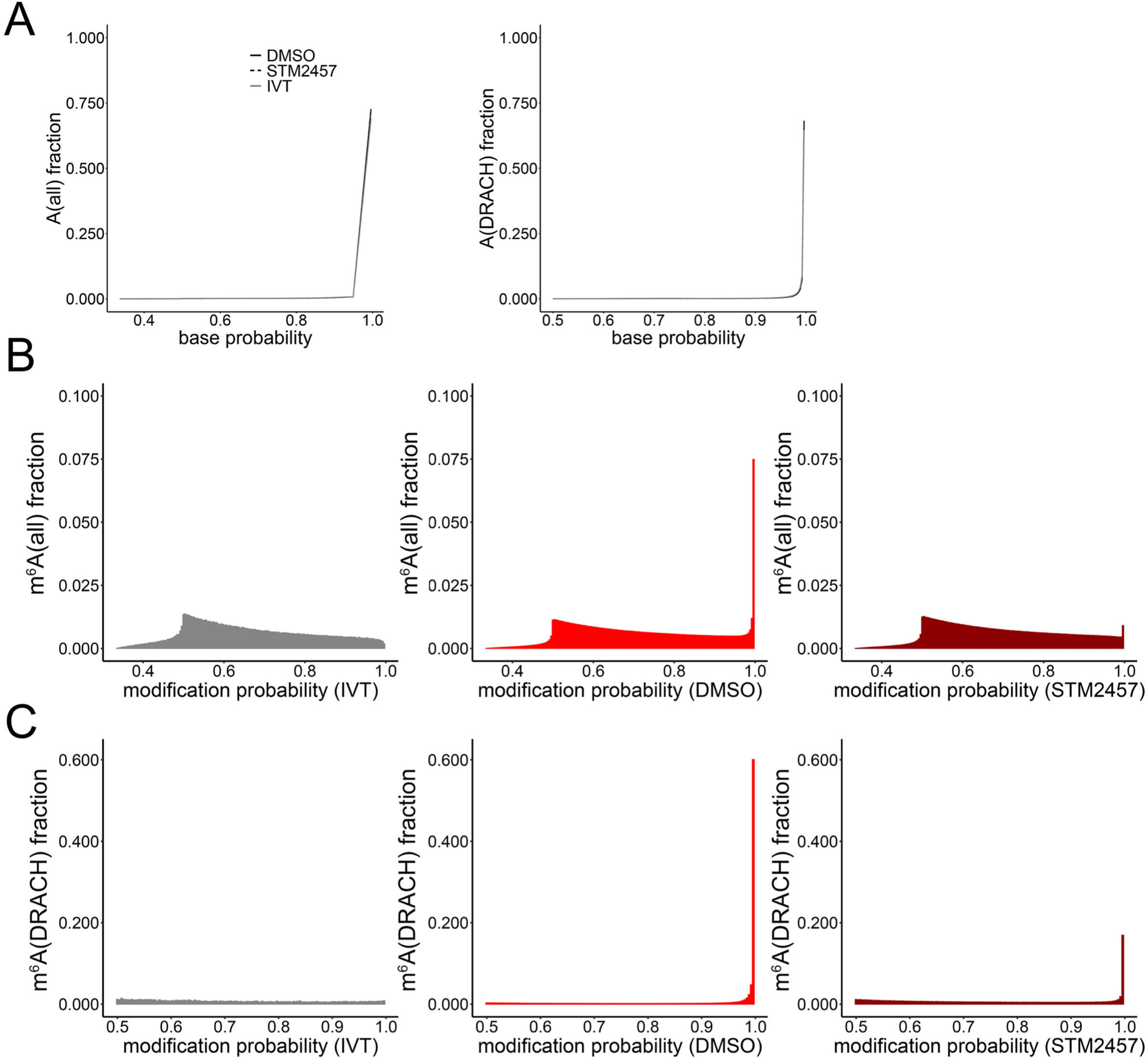
Evaluation of Dorado by base and modification probability distributions. **(A)** Dorado v0.9.0-generated basecall accuracy probability values for A (all-context and DRACH-context basecaller models) from genome-aligned DMSO-and STM2457-treated NHDF samples against IVT poly(A) NHDF RNA. A probability score distribution is shown as a black solid line for DMSO, STM2457 – black dashed line, IVT – grey solid line. **(B)** Dorado v0.9.0-generated modification probability distributions, visualized as histograms, for all-context m^6^A from genome-aligned DMSO-and STM2457-treated NHDFs and IVT NHDFs. IVT modification probability score distribution is shown in grey, DMSO – bright red, STM2457 – dark red. **(C)** Same as in (B), but for DRACH-context m^6^A sites.

**Figure S3.**
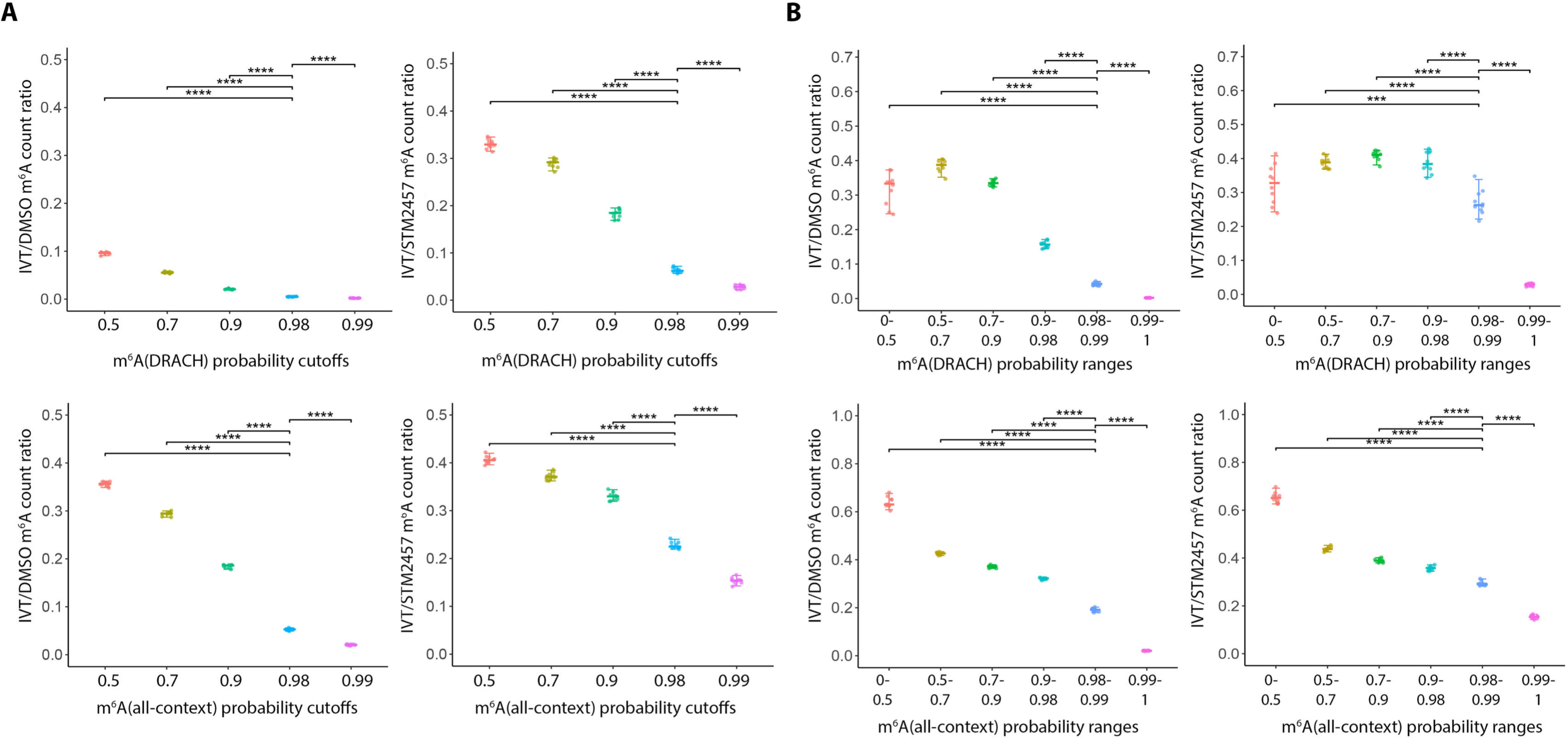
IVT-based estimation of background signal-derived m^6^A detection across modification probability thresholds and ranges in NHDFs. **(A)** m^6^A ratiosb derived from background signal across increasing modification probability cutoffs. For each cutoff (0.5, 0.7, 0.9, 0.98, and 0.99), the number of m^6^A calls detected in the IVT control was divided by the corresponding number detected in DMSO-or STM2457-treated NHDF samples. **(B)** m^6^A ratios derived from background signal calculated for discrete modification probability ranges (0–0.5, 0.5–0.7, 0.7–0.9, 0.9–0.98, 0.98–0.99, and 0.99–1). In both panels, the upper row shows DRACH-context m^6^A calls and the lower row shows all-context m^6^A calls; left and right plots show ratios relative to DMSO-and STM2457-treated samples, respectively. Each point represents an independently generated subsample of 200,000 reads. Statistical significance was assessed using pairwise comparisons derived from linear mixed-effects models, with sample included as a random effect and Benjamini–Hochberg correction for multiple comparisons. ****P < 0.0001.

**Figure S4.**
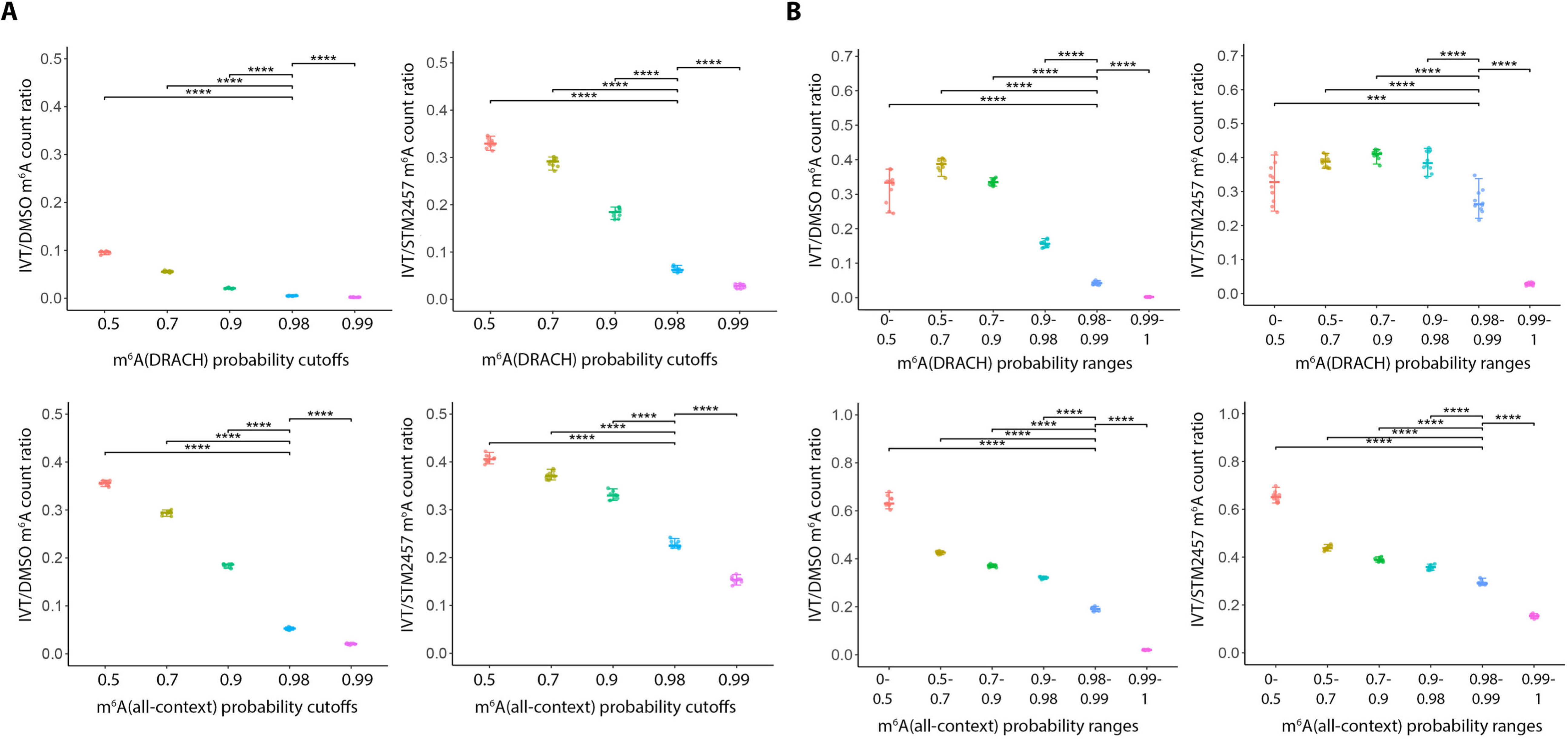
IVT-based estimation of m^6^A detection derived from background signal across modification probability thresholds and ranges in HD10.6 cells. **(A)** m^6^A ratios derived from background signal across increasing modification probability cutoffs. For each cutoff (0.5, 0.7, 0.9, 0.98, and 0.99), the number of m^6^A calls detected in the IVT control was divided by the corresponding number detected in DMSO-or STM2457-treated NHDF samples. **(B)** m^6^A ratios derived from background signal calculated for discrete modification probability ranges (0–0.5, 0.5–0.7, 0.7–0.9, 0.9–0.98, 0.98–0.99, and 0.99–1). In both panels, the upper row shows DRACH-context m^6^A calls and the lower row shows all-context m^6^A calls; left and right plots show ratios relative to DMSO-and STM2457-treated samples, respectively. Each point represents an independently generated subsample of 200,000 reads. Statistical significance was assessed using pairwise comparisons derived from linear mixed-effects models, with sample included as a random effect and Benjamini–Hochberg correction for multiple comparisons. ****P < 0.0001.

**Figure S5.**
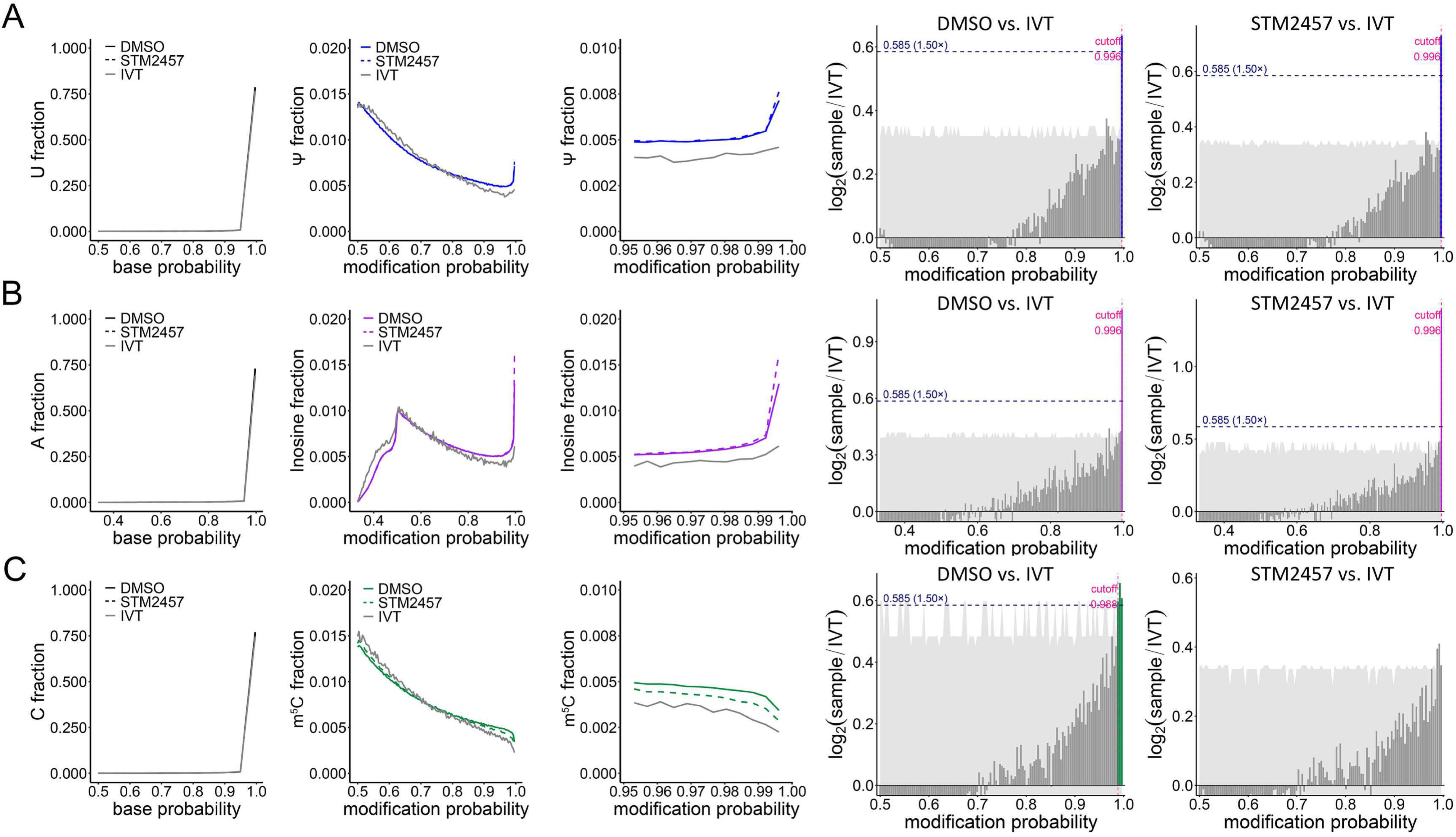
Modification probability cutoffs are necessary to identify Ψ and inosine sites more accurately. **(A)** (left) Dorado v0.9.0-generated basecall accuracy probability values for U (left) and modification probability distributions for Ψ (right) from genome-aligned DMSO-and STM2457-treated NHDF samples are plotted against IVT poly(A) RNA. Ψ probabilities are visualized in blue solid and dashed lines for DMSO and STM2457, respectively and in grey solid line for IVT. (right) Bar plots showing the per-bin log2(sample/IVT) ratio of modification probability fractions for the DMSO-and STM2457-treated NHDFs conditions relative to the unmodified IVT reference. The horizontal navy dashed line marks the fold-enrichment threshold used to call enrichment. Bars are coloured by significance relative to the fold-enrichment threshold (blue, log2 ≥ 1.5-fold; dark grey, not significant). The light grey ribbon indicates the 95% bootstrap confidence interval around the log2 ratio estimate at each bin. The vertical pink dashed line indicates the derived modification probability cutoff above which sites are considered genuinely modified. **(B)** Same as in (A) but showing A base accuracy probabilities and inosine sites. Inosine probabilities are visualized in purple solid and dashed lines for DMSO and STM2457, respectively and in grey solid line for IVT. **(C)** Same as in (A) but showing C base accuracy probabilities and m^5^C sites. m^5^C probabilities are visualized in green solid and dashed lines for DMSO and STM2457, respectively and in grey solid line for IVT.

**Figure S6.**
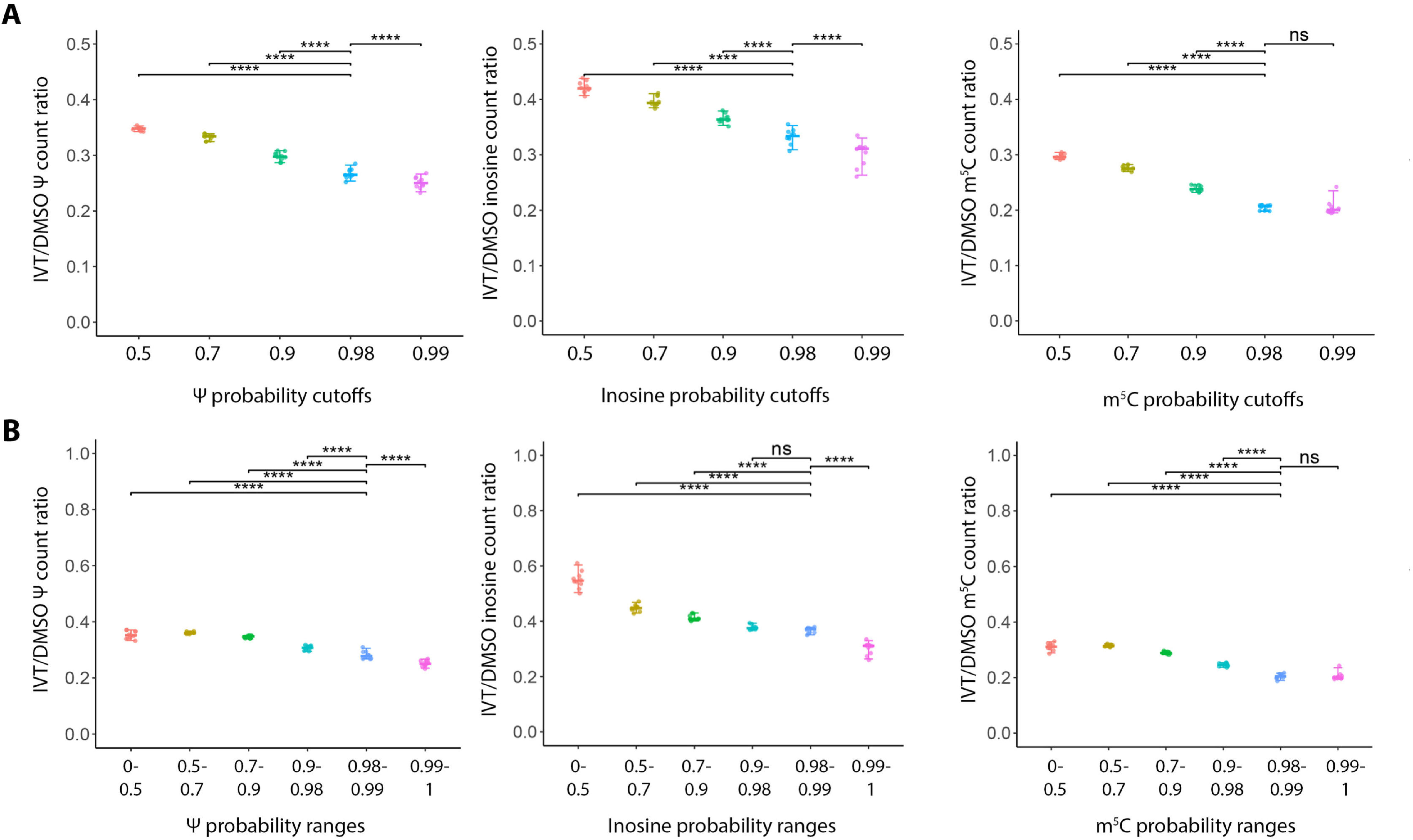
IVT-based estimation of Ψ, inosine, and m^5^C detection derived from background signal across modification probability thresholds and ranges in NHDFs. **(A)** Modification ratios derived from background signal across increasing modification probability cutoffs (0.5, 0.7, 0.9, 0.98, and 0.99) for Ψ, inosine, and m^5^C. For each cutoff, the number of modification calls detected in the IVT control was divided by the corresponding number detected in DMSO-treated NHDF samples. **(B)** Modification ratios derived from background signal calculated for discrete modification probability ranges (0–0.5, 0.5–0.7, 0.7–0.9, 0.9–0.98, 0.98–0.99, and 0.99–1). From left to right, plots show results for Ψ, inosine, and m^5^C. Each point represents an independently generated subsample of 200,000 reads. Statistical significance was assessed using pairwise comparisons derived from linear mixed-effects models, with sample included as a random effect and Benjamini–Hochberg correction for multiple comparisons. ****P < 0.0001; ns, not significant.

**Figure S7.**
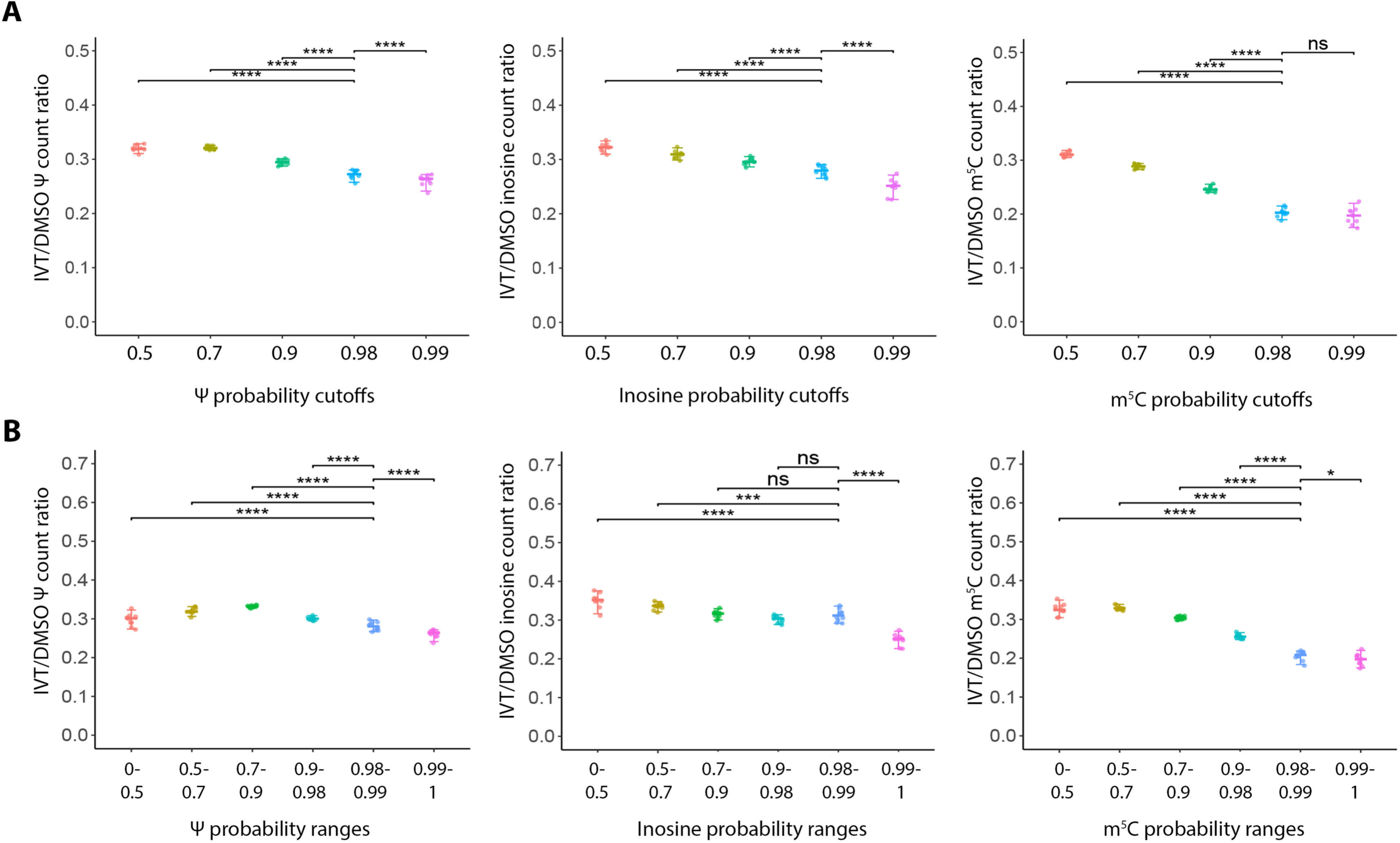
IVT-based estimation of Ψ, inosine, and m^5^C derived from background signal detection across modification probability thresholds and ranges in HD10.6 cells. **(A)** Modification ratios derived from background signal across increasing modification probability cutoffs (0.5, 0.7, 0.9, 0.98, and 0.99) for Ψ, inosine, and m^5^C. For each cutoff, the number of modification calls detected in the IVT control was divided by the corresponding number detected in DMSO-treated HD10.6 samples. **(B)** Modification ratios derived from background signal calculated for discrete modification probability ranges (0–0.5, 0.5–0.7, 0.7–0.9, 0.9–0.98, 0.98–0.99, and 0.99–1). From left to right, plots show results for Ψ, inosine, and m^5^C. Each point represents an independently generated subsample of 200,000 reads. Statistical significance was assessed using pairwise comparisons derived from linear mixed-effects models, with sample included as a random effect and Benjamini–Hochberg correction for multiple comparisons. ****P < 0.0001; ns, not significant.

**Figure S8.**
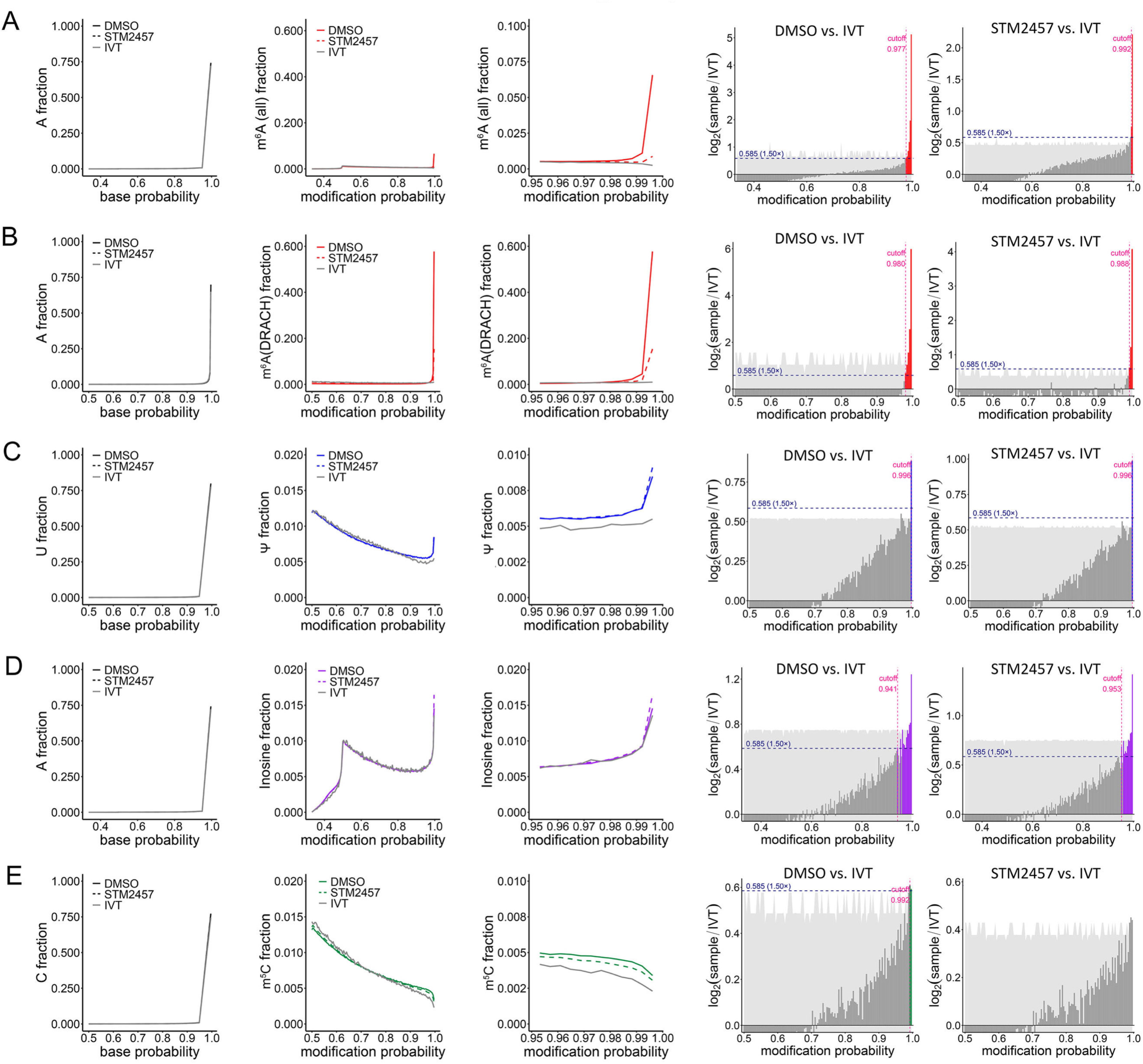
Modification probability cutoffs are necessary to identify modification sites accurately, shown for transcriptome-aligned data. Dorado v0.9.0-generated basecall accuracy probability values for A, U and C (left, **A-E**) and modification probability distributions for m^6^A (all-context), m^6^A (DRACH-context), Ψ, inosine and m^5^C (middle, **A-E**) in transcriptome-aligned DMSO-and STM2457-treated NHDFs, plotted against the values from IVT NHDF poly(A) RNA. (right) Bar plots showing the per-bin log2(sample/IVT) ratio of modification probability fractions for the DMSO-and STM2457-treated NHDFs conditions relative to the unmodified IVT reference. The horizontal navy dashed line marks the fold-enrichment threshold used to call enrichment. Bars are coloured by significance relative to the fold-enrichment threshold (blue, log2 ≥ 1.5-fold; dark grey, not significant). The light grey ribbon indicates the 95% bootstrap confidence interval around the log2 ratio estimate at each bin. The vertical pink dashed line indicates the derived modification probability cutoff above which sites are considered genuinely modified.

**Figure S9.**
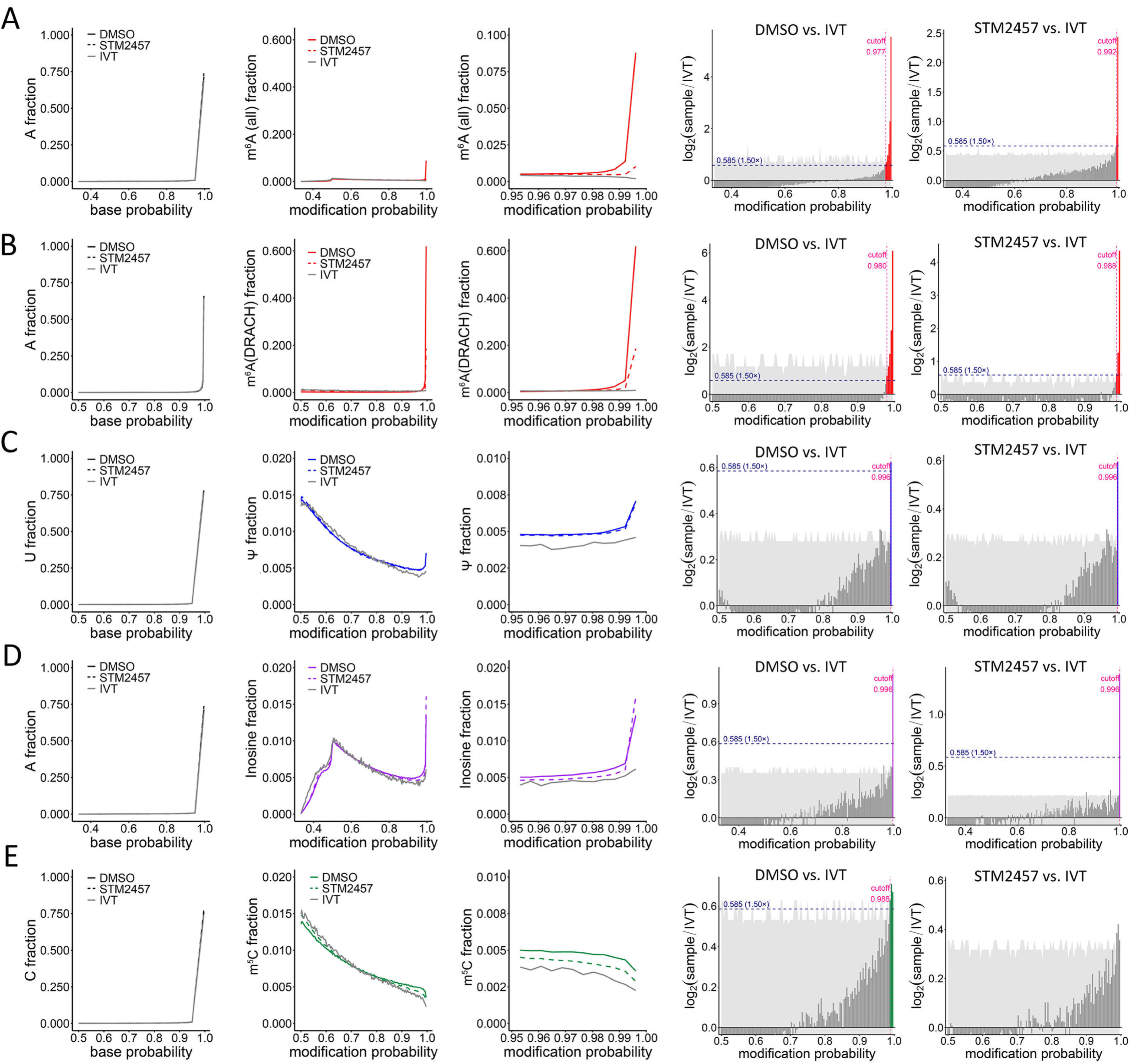
Modification probability distribution profiles in genome-aligned DMSO-and STM2457-treated HD10.6 cells. Dorado v0.9.0-generated basecall accuracy probability values for A, U and C (left, **A-E**) and modification probability distributions for m^6^A (all-context), m^6^A (DRACH-context), Ψ, inosine and m^5^C (middle, **A-E**) in human genome-aligned DMSO-and STM2457-treated HD10.6 cells, plotted against the values from IVT HD10.6 poly(A) RNA. (right) Bar plots showing the per-bin log2(sample/IVT) ratio of modification probability fractions for the DMSO-and STM2457-treated HD10.6 conditions relative to the unmodified IVT reference. The horizontal navy dashed line marks the fold-enrichment threshold used to call enrichment. Bars are coloured by significance relative to the fold-enrichment threshold (blue, log2 ≥ 1.5-fold; dark grey, not significant). The light grey ribbon indicates the 95% bootstrap confidence interval around the log2 ratio estimate at each bin. The vertical pink dashed line indicates the derived modification probability cutoff above which sites are considered genuinely modified.

**Figure S10.**
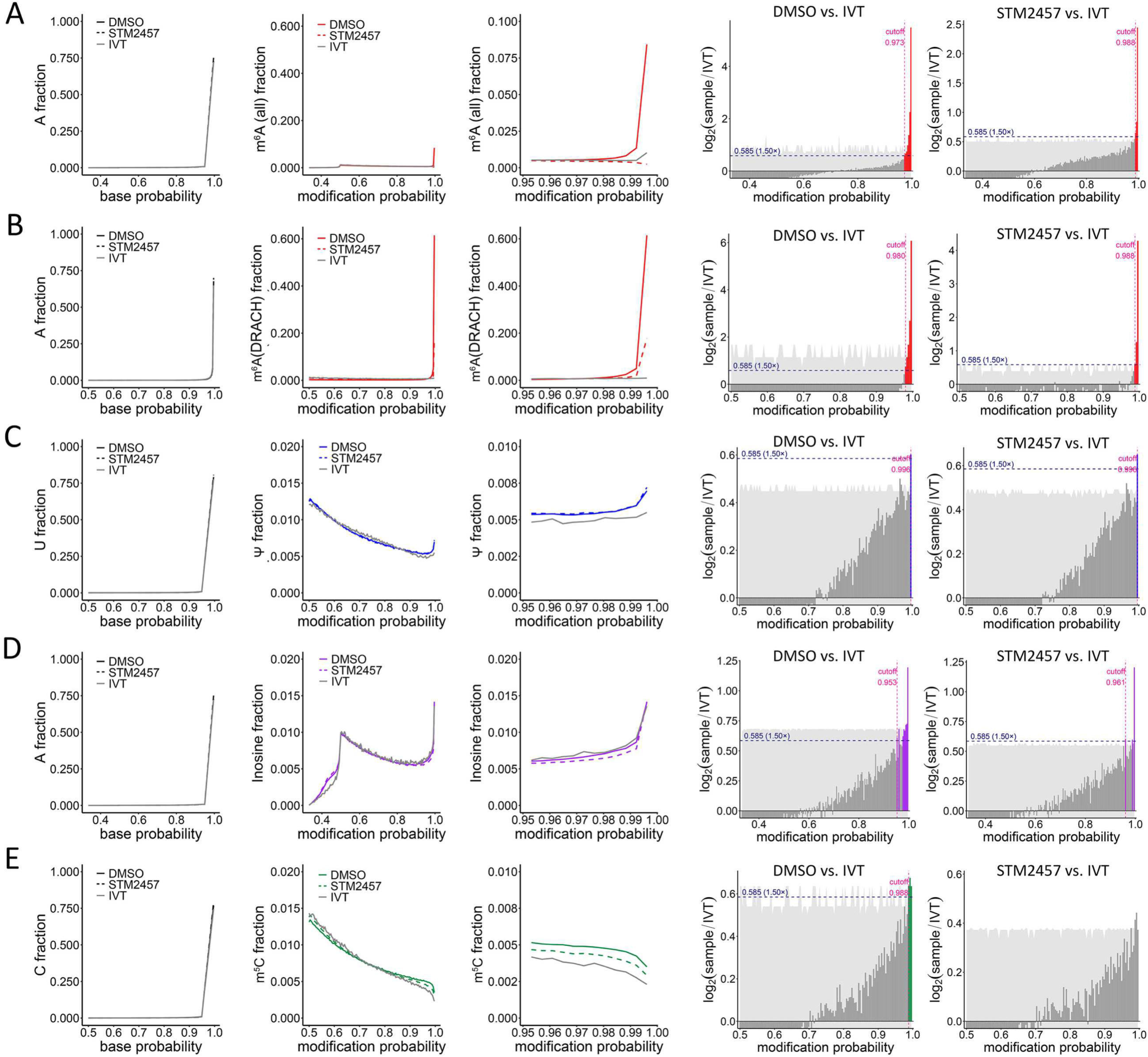
Modification probability distribution profiles in transcriptome-aligned DMSO-and STM2457-treated HD10.6 cells. Dorado v0.9.0-generated basecall accuracy probability values for A, U and C (left, **A-E**) and modification probability distributions for m^6^A (all-context), m^6^A (DRACH-context), Ψ, inosine and m^5^C (middle, **A-E**) in human transcriptome-aligned DMSO-and STM2457-treated HD10.6 cells, plotted against the values from IVT HD10.6 poly(A) RNA. (right) Bar plots showing the per-bin log2(sample/IVT) ratio of modification probability fractions for the DMSO-and STM2457-treated HD10.6 conditions relative to the unmodified IVT reference. The horizontal navy dashed line marks the fold-enrichment threshold used to call enrichment. Bars are coloured by significance relative to the fold-enrichment threshold (blue, log2 ≥ 1.5-fold; dark grey, not significant). The light grey ribbon indicates the 95% bootstrap confidence interval around the log2 ratio estimate at each bin. The vertical pink dashed line indicates the derived modification probability cutoff above which sites are considered genuinely modified.

**Figure S11.**
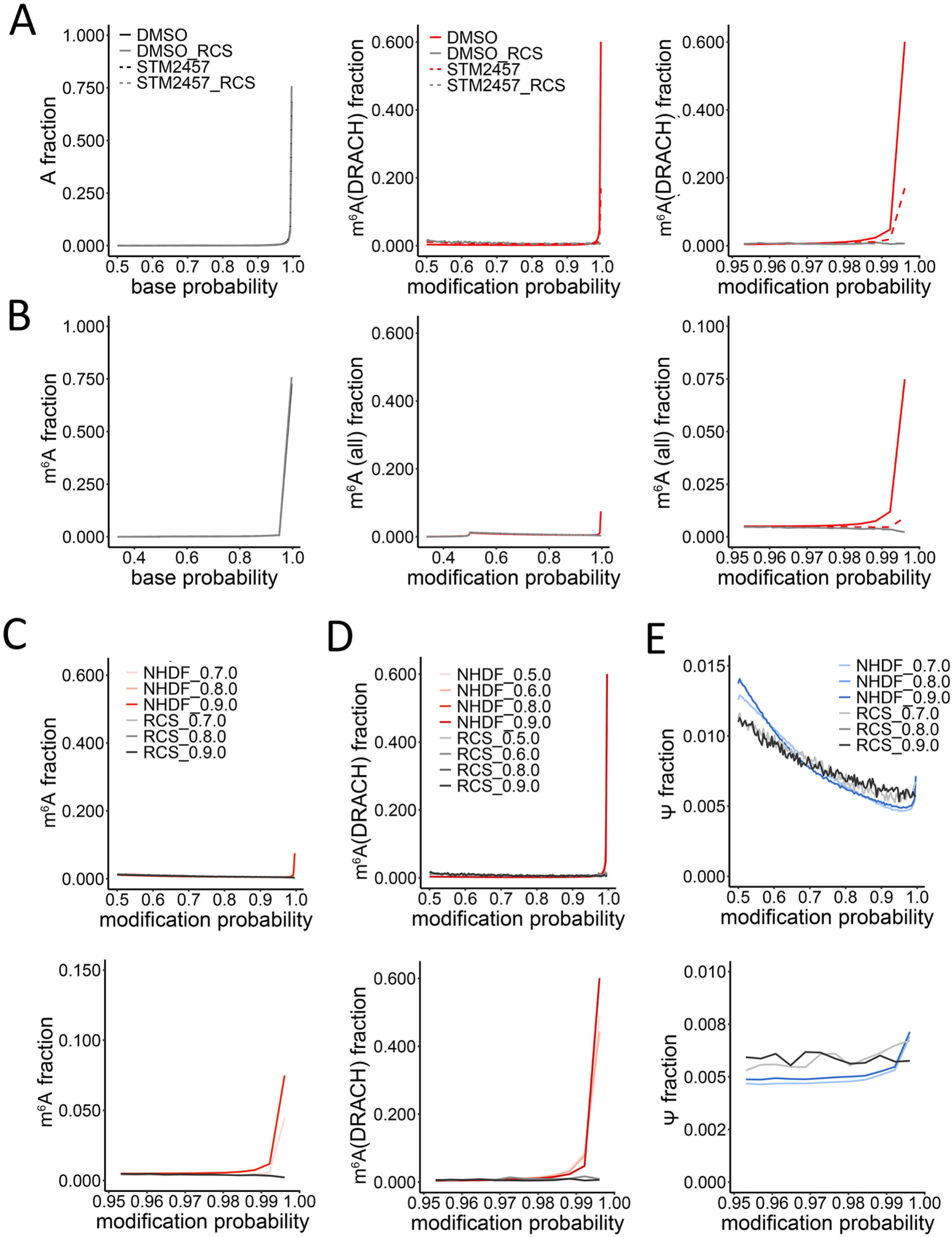
Modification site probabilities detected with different versions and models of Dorado and compared to RCS. **(A,B)** DRACH-context and all-context m^6^A modification probability distributions from genome-aligned DMSO (solid red line)-and STM2457 (dashed red line)-treated NHDFs are plotted against RCS negative control (grey). **(C)** Modification probability distributions for all-context m^6^A (Dorado v0.7.0 (model v5.0.0), v0.8.0 (model v5.1.0) and v0.9.0 (model v5.1.0)), the bottom panel showing the distributions in 0.95-1.00 modification range. **(D)** Same as in (C), but for DRACH-context m^6^A (Dorado v0.5.0 (model v3.0.1), v0.6.0 (model v3.0.1), v0.8.0 (model v5.1.0) and v0.9.0 (model v5.1.0)). **(E)** Same as in (C), but for Ψ (Dorado v0.7.0 (model v5.0.0), v0.8.0 (model v5.1.0), v0.9.0 (model v5.1.0)). All the plots in C-E are generated from genome-aligned DMSO-treated NHDF datasets and are plotted against RCS negative controls.

**Figure S12.**
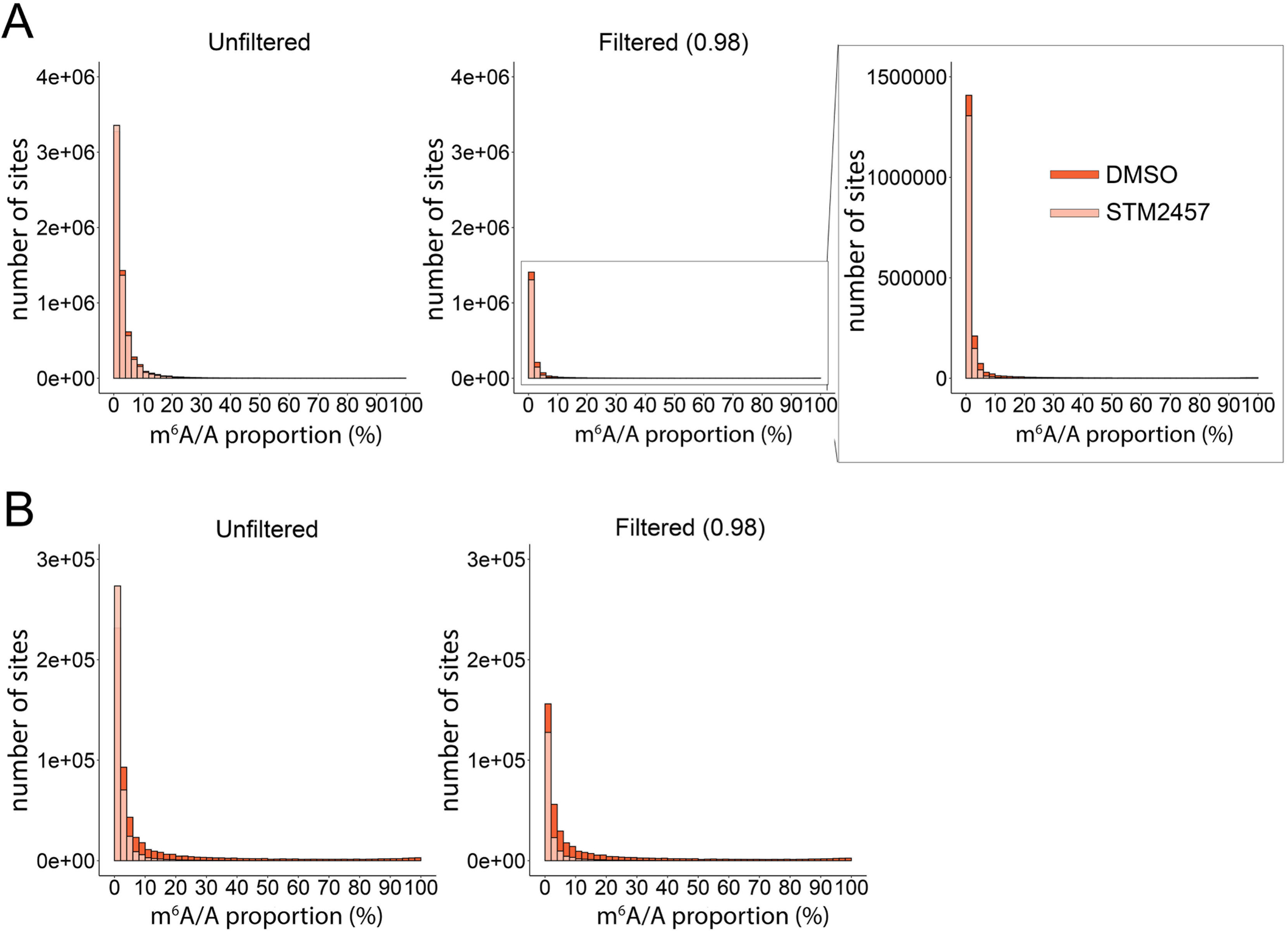
Evaluation of Dorado by the distributions of predicted m^6^A/A proportions. **(A)** Distributions of predicted m^6^A/A proportions for all-context m^6^A from DMSO-(in red) or STM2457-treated (in pink) NHDF datasets (genome-aligned and filtered for ≥ 20 reads) are plotted, comparing 0.98 modification probability-unfiltered (left) and filtered (right) sites. **(B)** Same as in (A), but for DRACH-context m^6^A sites.

**Figure S13.**
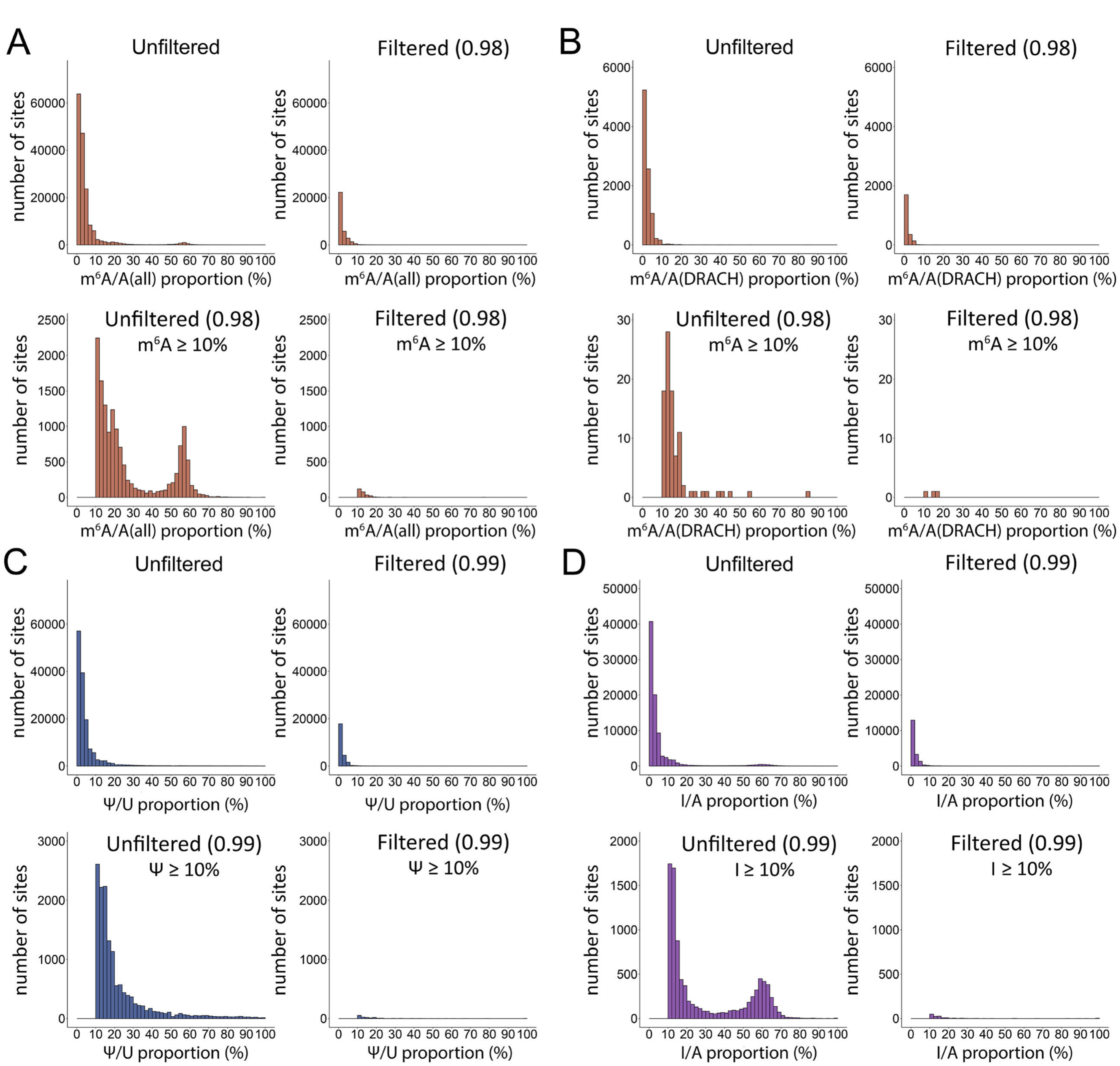
Distributions of predicted m^6^A/A, Ψ/U and inosine/A proportions in IVT datasets. **(A)** All-context predicted m^6^A/A proportion distributions (all proportions (top) and ≥ 10% proportions (bottom)) in (genome-aligned) IVT RNA from DMSO-treated NHDFs, comparing modification probability-unfiltered and filtered data. The bottom panel shows distributions only for ≥ 10% proportion sites. **(B)** As in (A), but for DRACH-context m^6^A. **(C)** As in (A), but for Ψ. **(D)** As in (A), but for inosine.

**Figure S14.**
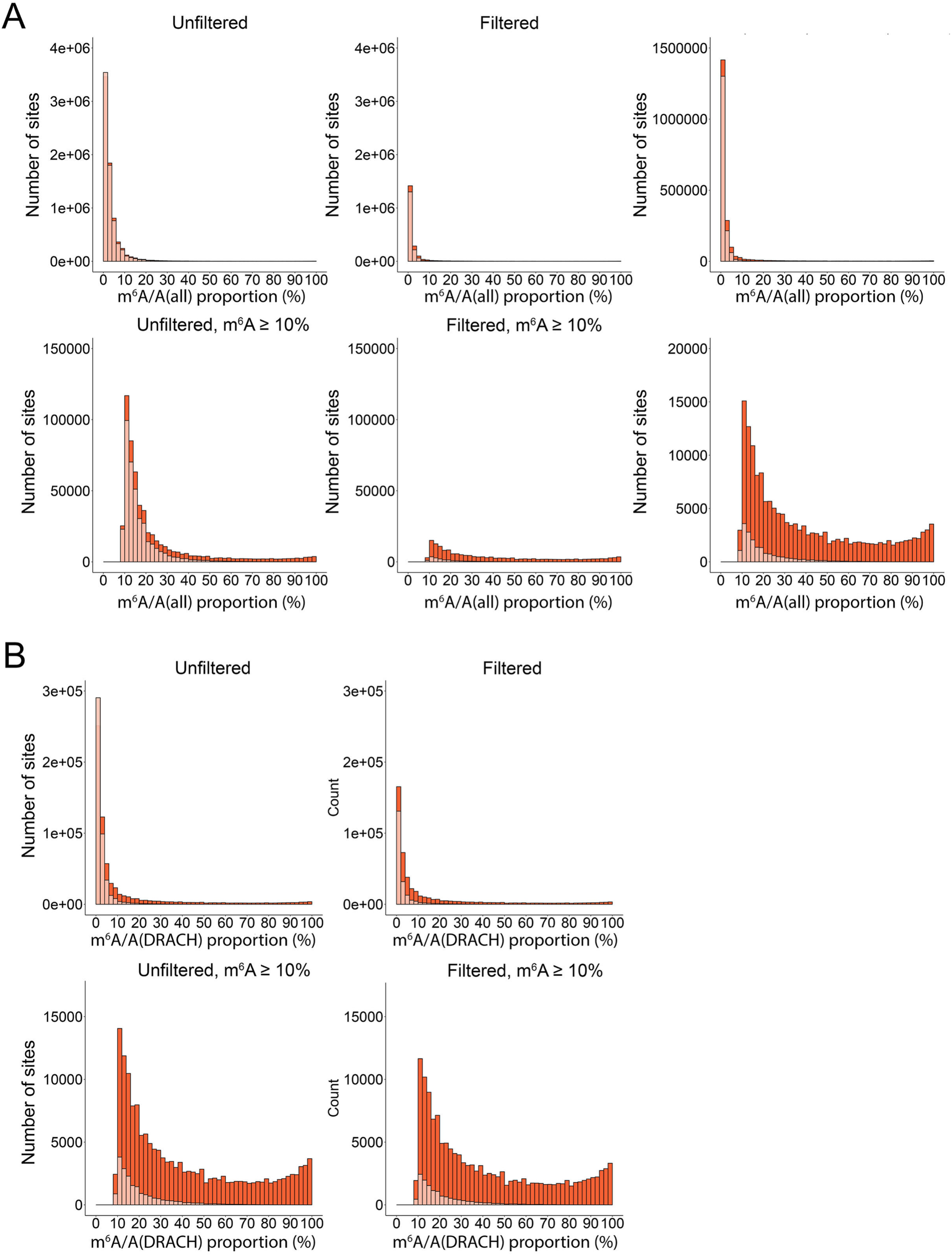
Evaluation of Dorado by predicted modification proportion distributions in transcriptome-aligned NHDFs. **(A)** Distributions of predicted m^6^A/A proportions for all-context m^6^A from DMSO-(in red) or STM2457-treated (in pink) NHDF datasets (transcriptome-aligned and filtered for ≥ 20 reads) are plotted, assessing the 0.98 modification probability cutoff. The bottom panel shows distributions only for ≥ 10%-proportion sites. **(B)** Same as in (A), but for DRACH-context m^6^A sites.

**Figure S15.**
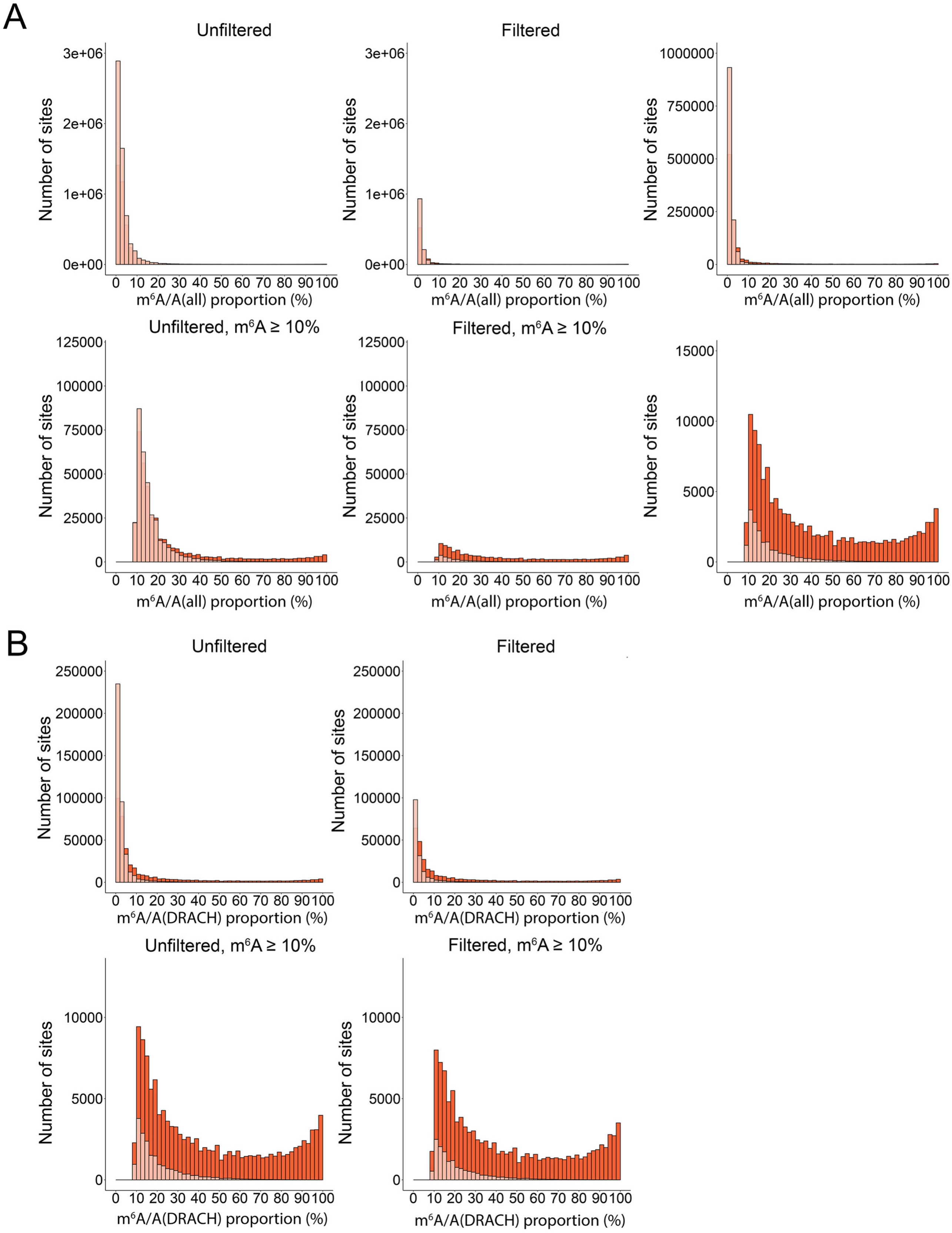
Evaluation of Dorado by distributions of predicted m^6^A/A proportions in transcriptome-aligned HD10.6 cells. **(A)** Distributions of predicted m^6^A/A proportions for all-context m^6^A from DMSO-(in red) or STM2457-treated (in pink) HD10.6 datasets (transcriptome-aligned and filtered for ≥ 20 reads) are plotted, assessing the 0.98 modification probability cutoff. The bottom panel shows distributions only for ≥ 10%-proportion sites. **(B)** Same as in (A), but for DRACH-context m^6^A sites.

**Figure S16.**
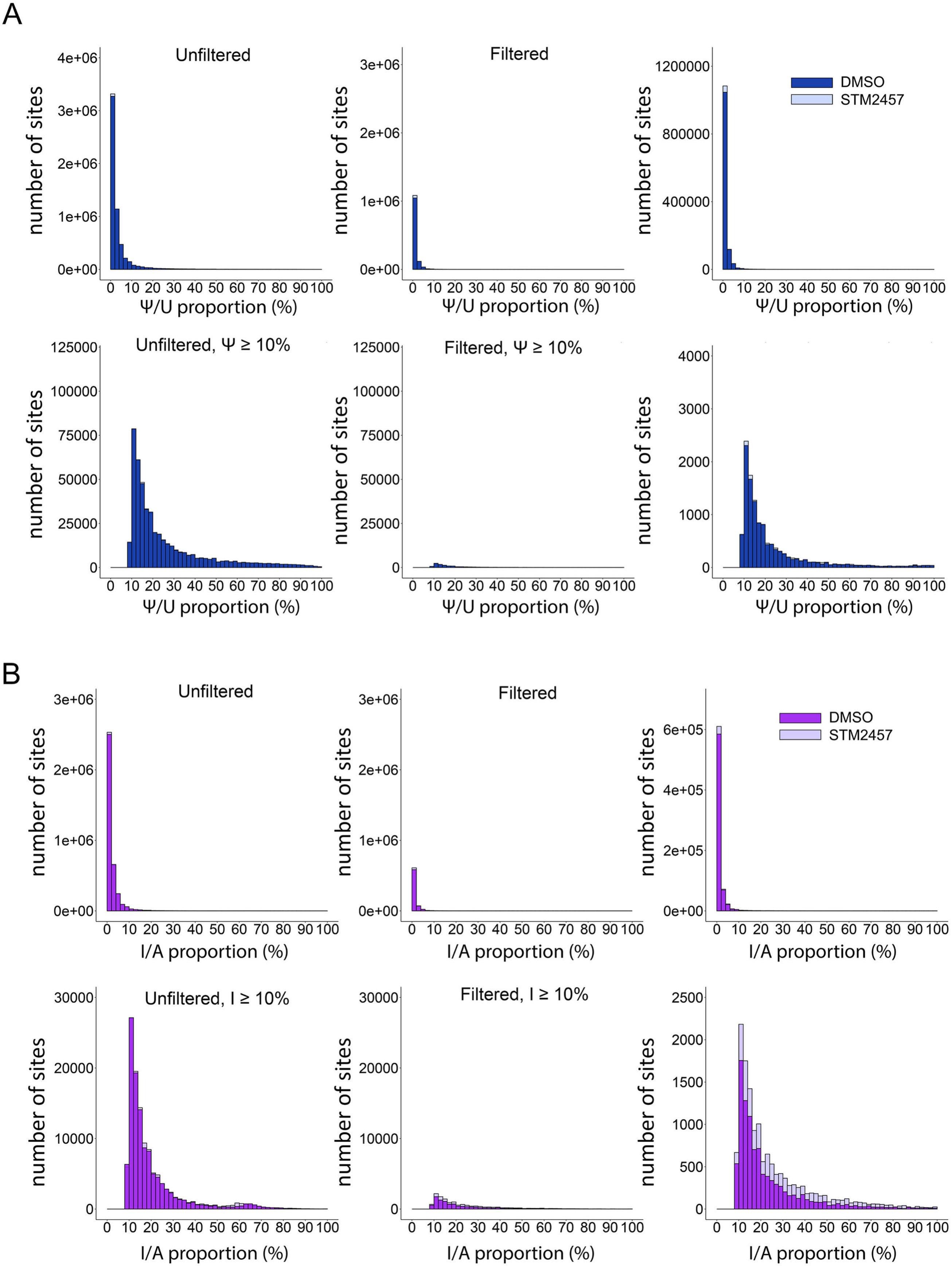
Distributions of predicted Ψ/U and inosine/A proportions in NHDFs. **(A)** Distributions of all (left) and ≥ 10% (right) predicted Ψ/U proportions in the 0.99 modification probability-unfiltered and filtered data from genome-aligned DMSO-(darker color) and STM2457-treated (lighter color) NHDFs (filtered for ≥ 20 reads). **(B)** Same as in (A), but for inosine.

**Figure S17.**
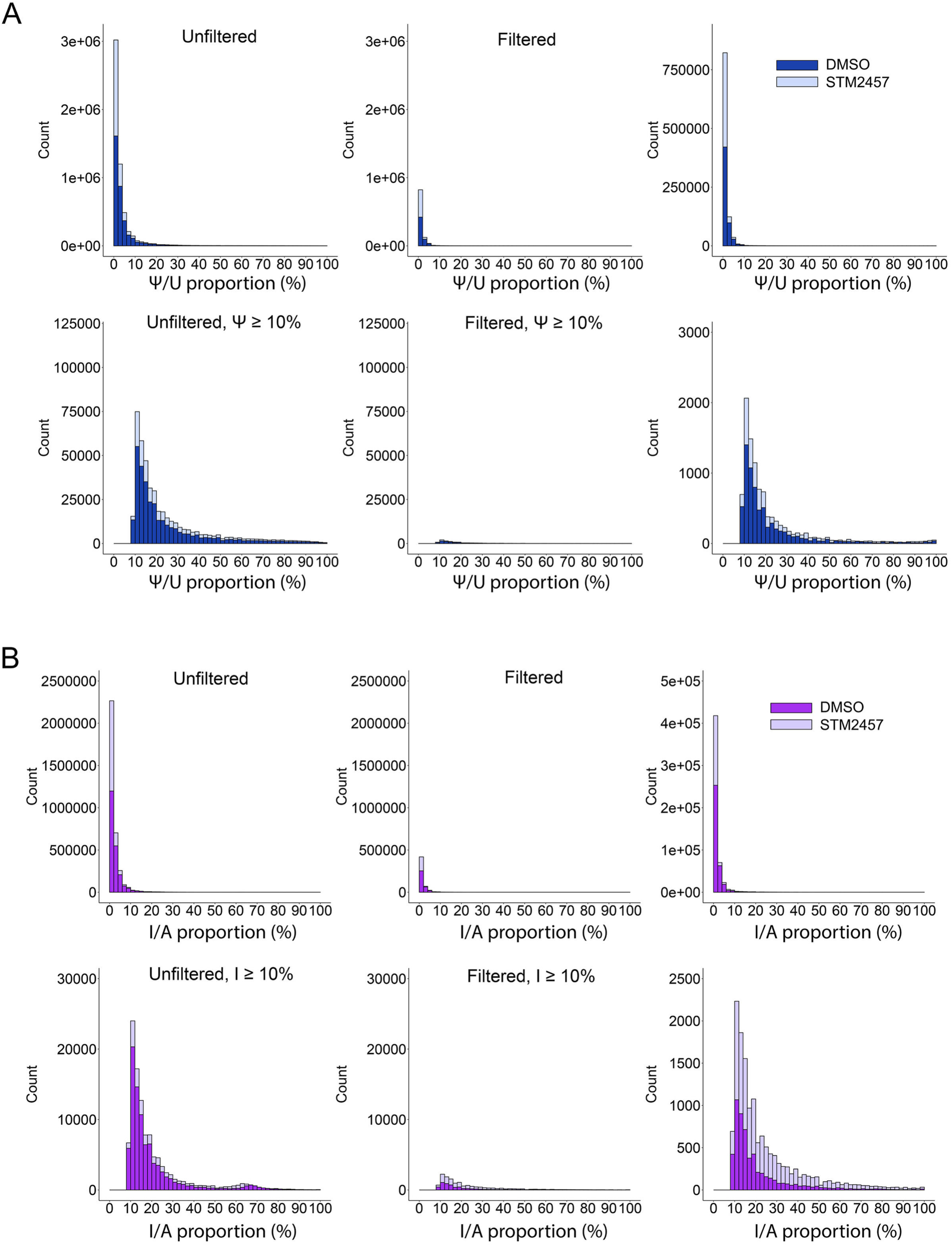
Distributions of predicted Ψ/U and inosine/A proportions in HD10.6 cells. **(A)** Distributions of all and ≥ 10% predicted Ψ/U proportions in the 0.99 modification probability-unfiltered and filtered data from genome-aligned DMSO-(darker color) and STM2457-treated (lighter color) HD10.6 cells (filtered for ≥ 20 reads). **(B)** Same as in (A), but for inosine.

**Figure S18.**
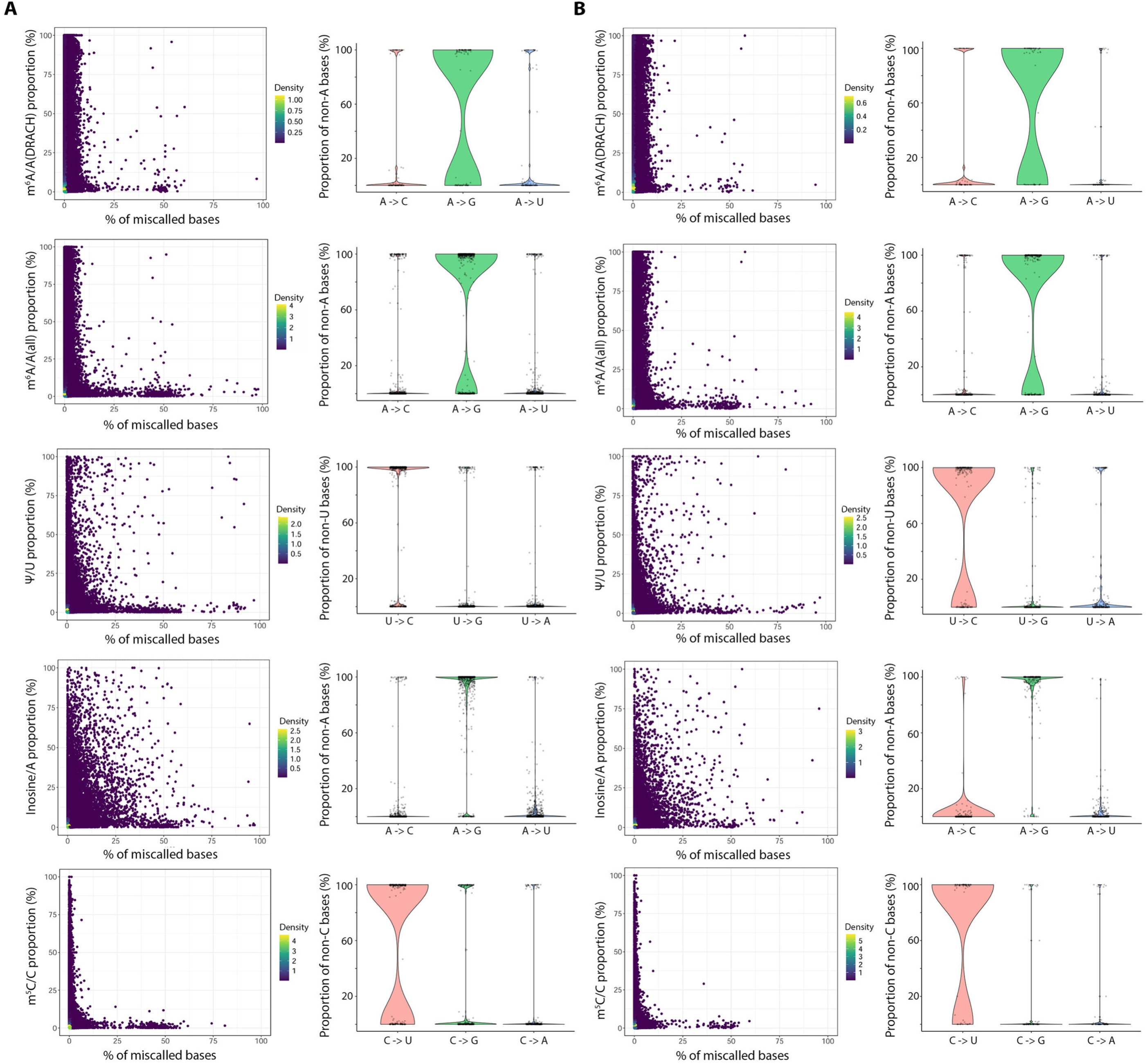
Relationship between predicted RNA modification proportions and nucleotide basecalling errors in NHDF and HD10.6 datasets. **(A)** NHDF and **(B)** HD10.6 datasets showing, from top to bottom, DRACH-context m^6^A, all-context m^6^A, Ψ, inosine, and m^5^C sites. Prior to analysis, sites were filtered using modification probability thresholds of ≥0.98 for m^6^A and ≥0.99 for Ψ, inosine, and m^5^C, together with a minimum valid-read coverage of 20. For each modification type, scatterplots show the predicted modification proportion generated by Dorado and the percentage of overlapping reads basecalled as a nucleotide other than the corresponding reference nucleotide (A for m^6^A and inosine, T for Ψ, and C for m^5^C). Each point represents an individual modification site, with point color indicating density. Adjacent violin plots characterize the identity of alternative basecalls at sites for which ≥25% of all overlapping reads were basecalled as a non-reference nucleotide. For each site, the proportion of mis-basecalled reads assigned to each of the three alternative nucleotides was calculated, with individual points representing individual modification sites. Violin distributions therefore show the relative contribution of each alternative nucleotide among mis-basecalled reads at sites exhibiting high levels of basecalling mismatch.

**Figure S19.**
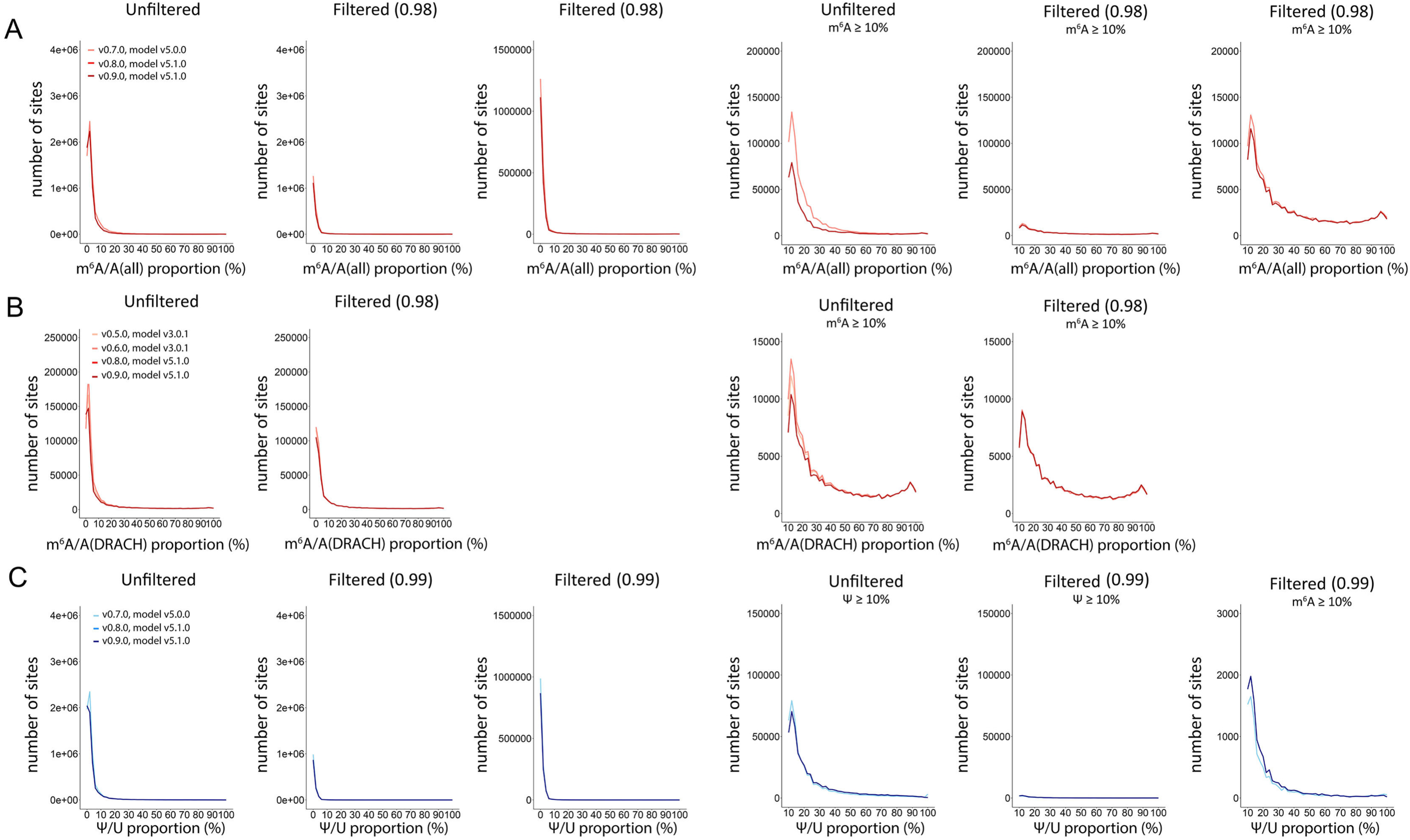
Predicted m^6^A/A and Ψ/U proportion distributions in different versions of Dorado. Distributions of predicted modification proportions (all – left, ≥ 10% – right) in a line graph format for **(A)** all-context m^6^A detected in genome-aligned DMSO-treated NHDF dataset, with Dorado versions 0.7.0 (model v5.0.0), 0.8.0 (model v5.1.0) and 0.9.0 (model v5.1.0) colored in gradients of red, darkness increasing with the later versions. **(B)** DRACH-context m^6^A detected with Dorado versions 0.5.0 (model v3.0.1), 0.6.0 (model v3.0.1), 0.8.0 (model v5.1.0) and 0.9.0 (model v5.1.0), colored in gradients of red, darkness increasing with the later versions. **(C)** Ψ detected with Dorado versions 0.7.0 (model v5.0.0), 0.8.0 (model v5.1.0) and 0.9.0 (model v5.1.0), colored in gradients of blue, darkness increasing with the later versions. All the datasets were filtered for ≥ 20 reads.

**Figure S20.**
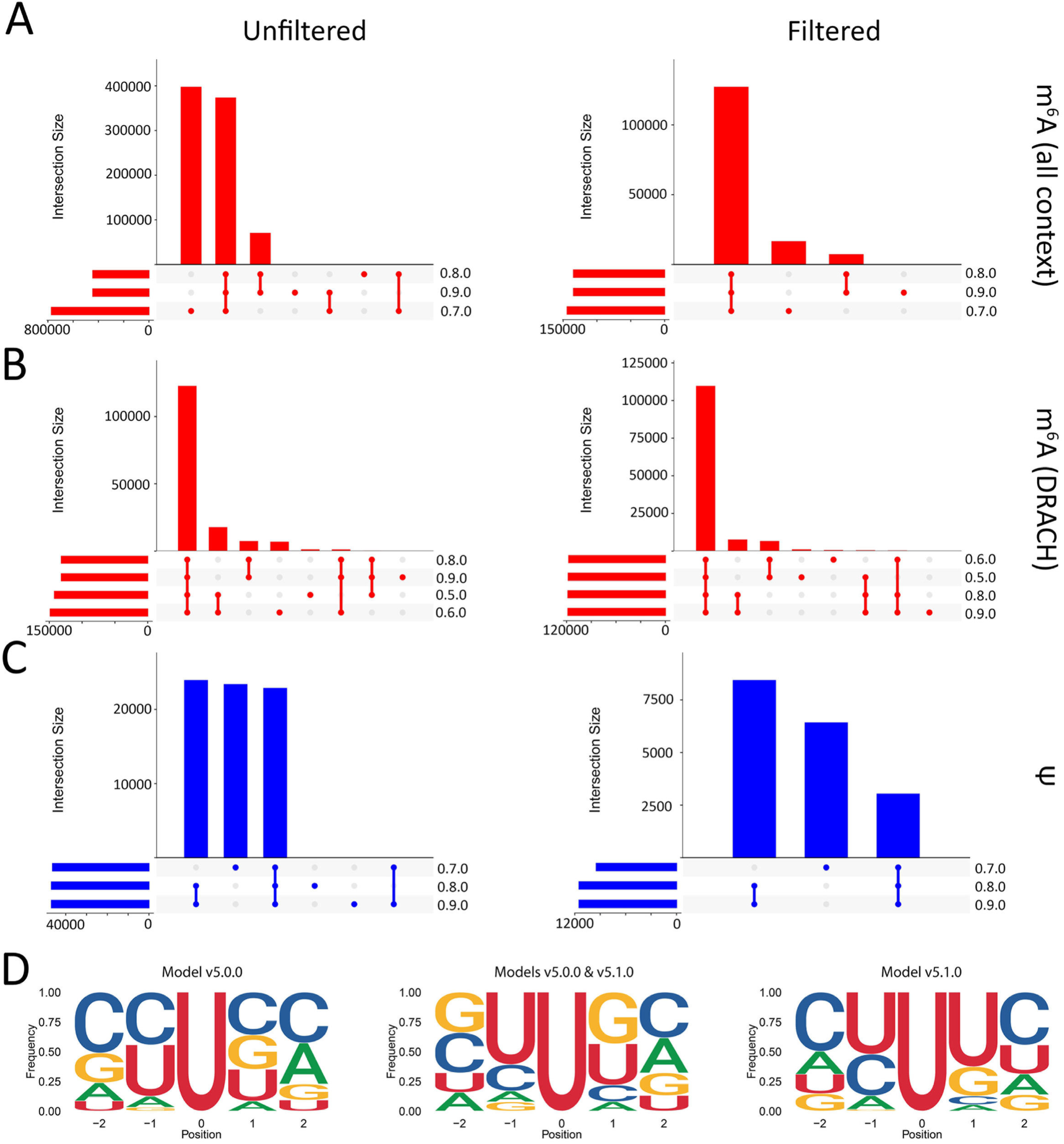
Comparison of modification sites detected with different versions of Dorado. Upset plots showing unique and shared detected modifications (filtered for ≥ 20 reads, ≥ 10% predicted modification proportion) in modification probability-unfiltered and filtered genome-aligned DMSO-treated NHDF data. The versions are ordered on the plot by the number of detected sites, in ascending order (on the left). Results are shown for the following models and versions: **(A)** all-context m^6^A detected with v.0.7.0 (model v5.0.0), v0.8.0 (model v5.1.0) and v0.9.0 (model v5.1.0), **(B)** DRACH-context m^6^A detected with v.0.5.0 (model v3.0.1), v0.6.0 (model v3.0.1), v0.8.0 (model v5.1.0) and v0.9.0 (model v5.1.0), **(C)** Ψ detected with v.0.7.0 (model v5.0.0), v0.8.0 (model v5.1.0) and v0.9.0 (model v5.1.0). **(D)** Nucleotide frequencies in the 5-base motif of Ψ sites uniquely for model v5.0.0 (basecalled with Dorado v0.7.0), common for models v5.0.0 (basecalled with v0.7.0) and v5.1.0 (basecalled with Dorado v0.8.0 and v0.9.0), and uniquely for model v5.1.0-detected (basecalled with v0.8.0 and v0.9.0) Ψ sites. Position 0 marks the modification site.

**Figure S21.**
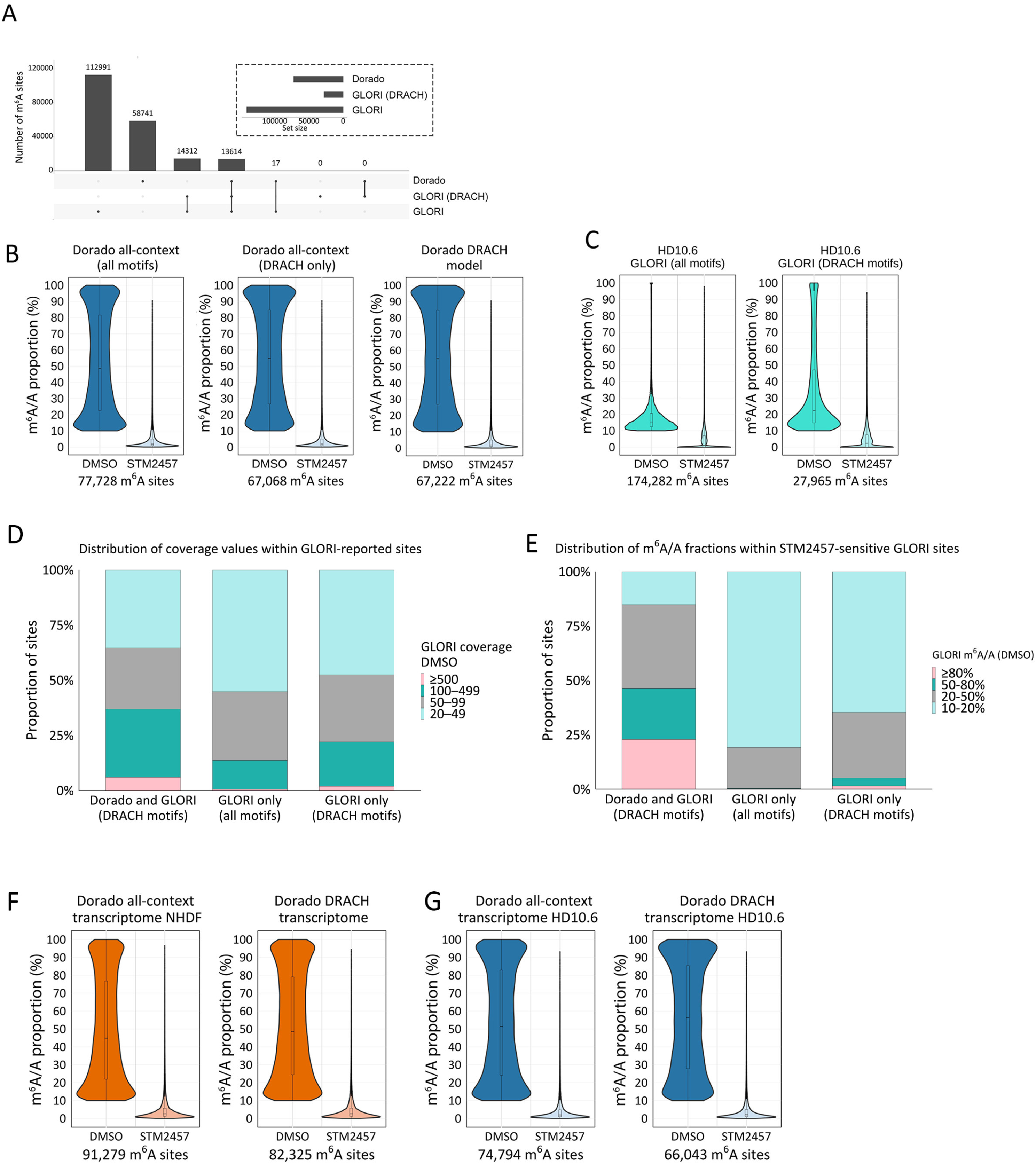
Comparison of m^6^A sites detected by Dorado and GLORI-Sequencing in NHDFs and HD10.6s. **(A)** Upset plot showing the number of m^6^A sites detected by GLORI and Dorado in NHDFs and their overlaps. Dorado sites represent high-confidence m^6^A sites obtained after filtering DRACH-model detected data. Similarly, only high confidence m^6^A sites detected by GLORI are shown but were additionally filtered for DRACH sites. **(B)** Distributions of predicted m^6^A/A proportions are plotted from genome-aligned DMSO-or STM2457-treated HD10.6 datasets using the indicated Dorado models. **(C)** Distributions of predicted m^6^A/A proportions from GLORI performed on DMSO-or STM2457-treated NHDFs are plotted. In addition, m^6^A sites were filtered for DRACH motifs (right plot). **(D)** Distribution of DMSO read coverage across STM2457-sensitive m^6^A sites reported in the GLORI dataset from NHDFs. Sites were categorized as those shared between GLORI and Dorado (DRACH motifs), GLORI-only across all motifs (non-DRACH sites), and GLORI-only sites within DRACH motifs. Within each category, sites were assigned to GLORI DMSO coverage bins (20-49, 5099, 100-499, and ≥ 500 reads), and the proportion of sites in each bin is shown on the y-axis. **(E)** Distribution of GLORI m^6^A/A stoichiometry among STM2457-sensitive m6A sites identified in NHDFs. Sites were categorized as shared between GLORI and Dorado within DRACH motifs, GLORI-only non-DRACH sites, and GLORI-only DRACH sites. Within each category, sites were assigned to GLORI DMSO m^6^A/A stoichiometry bins (10-20%, 20-50%, 50-80%, and ≥ 80%). Stacked bars show the proportion of sites in each stoichiometry bin. **(F)** Distributions of predicted m^6^A/A proportions are plotted from transcriptome-aligned DMSO-or STM2457-treated NHDF datasets using the indicated Dorado models. **(G)** Distributions of predicted m^6^A/A proportions are plotted from transcriptome-aligned DMSO-or STM2457-treated HD10.6 datasets using the indicated Dorado models.

**Figure S22.**
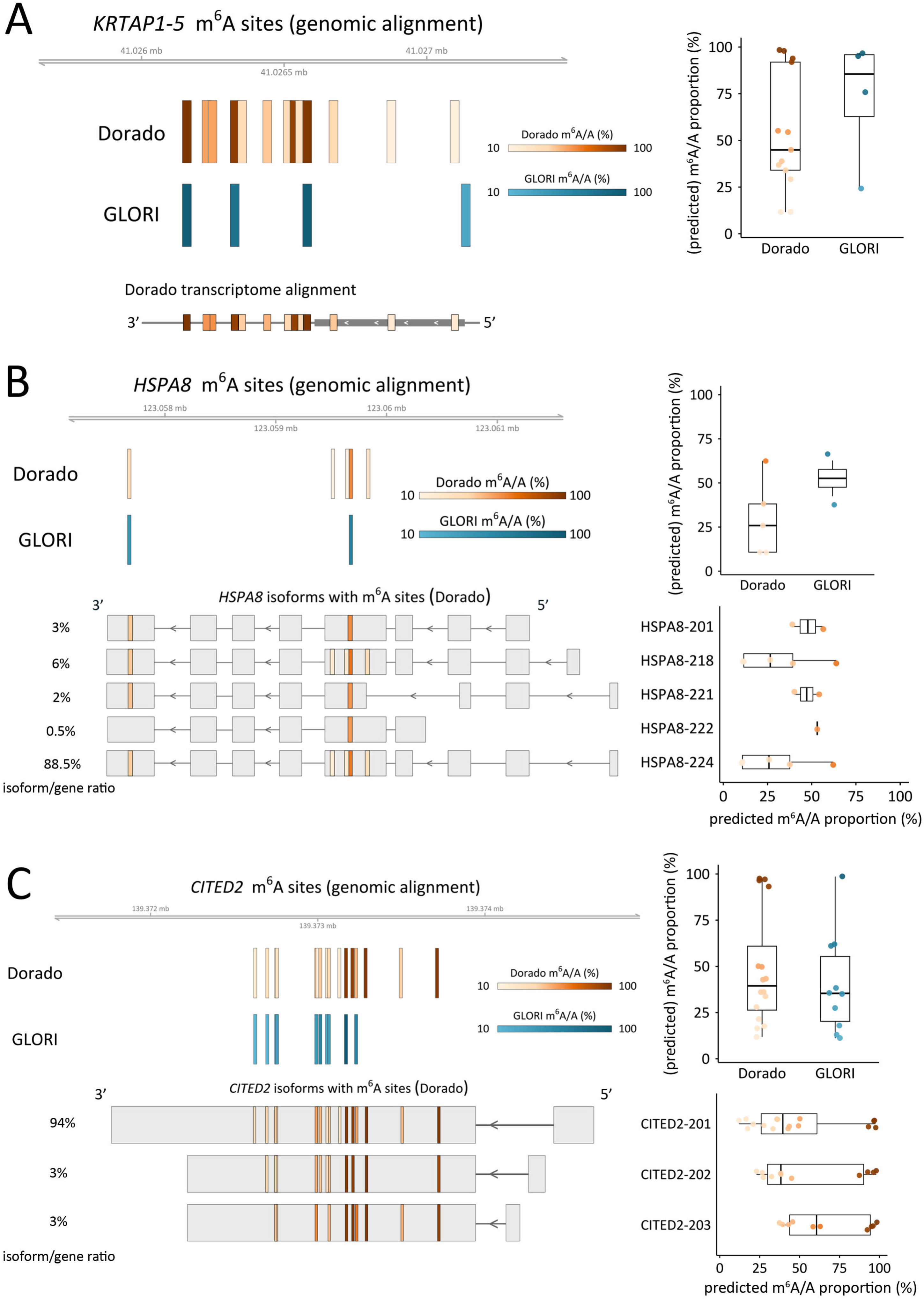
Isoform-level m^6^A prediction using the Dorado DRACH model in NHDF data. **(A)** m^6^A sites in the *KRTAP1-5* gene detected by GLORI and Dorado. Boxplots (right) represent the predicted m^6^A/A proportions per m^6^A site shown on the left. **(B)** m^6^A sites in the *HSPA8* gene detected by GLORI and Dorado. Boxplots (right) represent the predicted m^6^A/A proportions per m^6^A site shown on the left. Below, *HSPA8* isoforms with m^6^A sites, determined by Dorado. Boxplots (right) show the distribution of predicted m^6^A/A proportions on the m^6^A sites on *HSPA8-201* and *HSPA8-202*. **(C)** Same as in B, but for *CITED2*.

**Figure S23.**
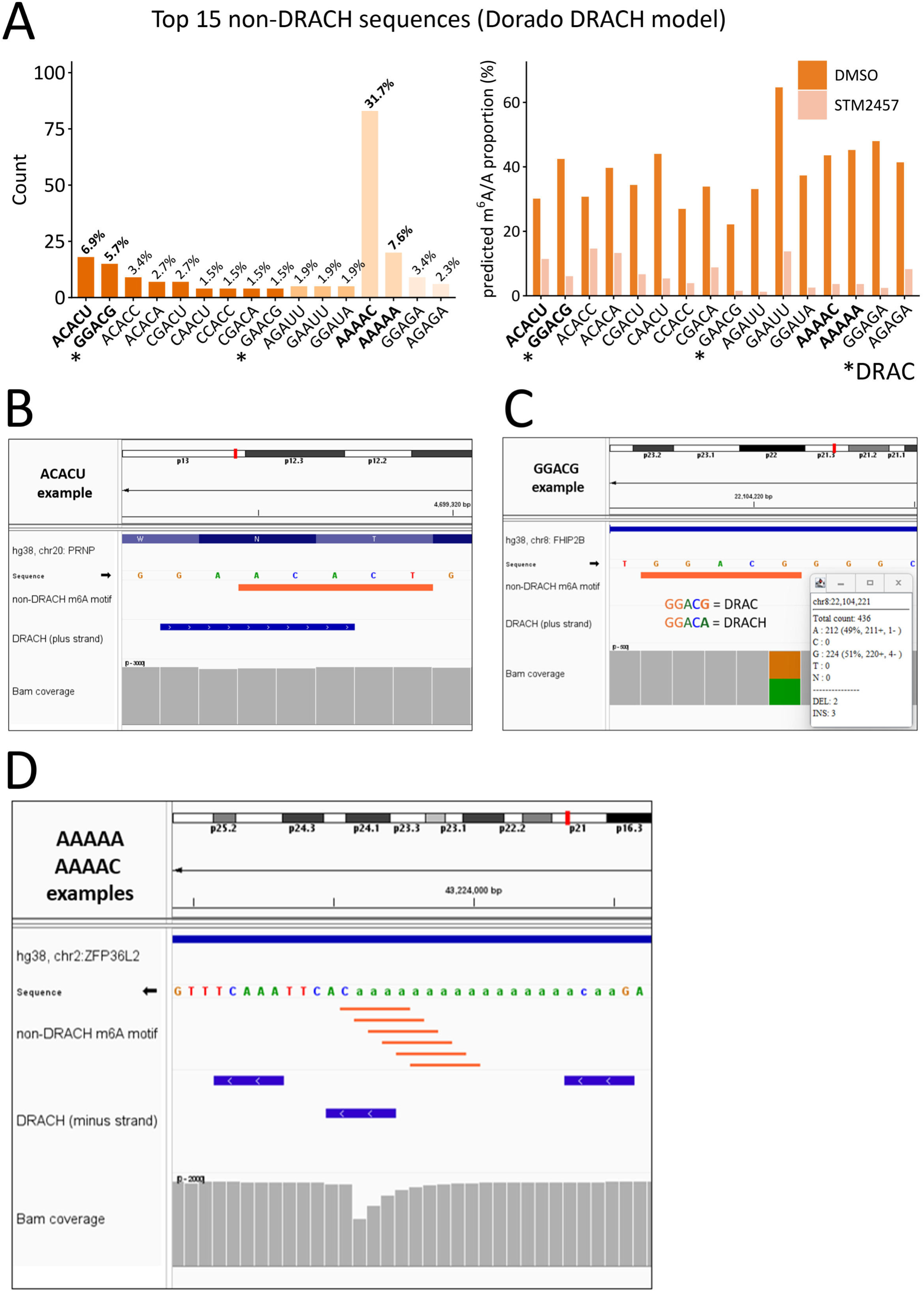
Non-DRACH m^6^A calls predicted by the Dorado DRACH model in NHDF data. **(A)** Top 15 most frequently detected non-DRACH motifs, ranked by occurrence (left); bars are colored by the central dinucleotide, and percentages indicate the proportion of all non-DRACH calls. Bold labels mark the 4 most abundant motifs discussed in the text. Mean predicted m**^6^**A/A proportions for each motif in DMSO-and STM2457-treated cells (right). Motifs conforming to DRAC but not DRACH are indicated with asterisks. **(B)** Representative genome browser view of an ACACU call located one to two nucleotides from a canonical DRACH motif, with the called and adjacent positions marked. **(C)** Read-level basecalls at a GGACG site, where the terminal position is called as G in 51% and A in 49% of reads, rendering the context non-DRACH and DRACH in approximately equal read fractions. **(D)** Representative AAAAA/AAAAC calls within a homopolymeric region. Neighboring DRACH motifs are indicated in blue.

**Figure S24.**
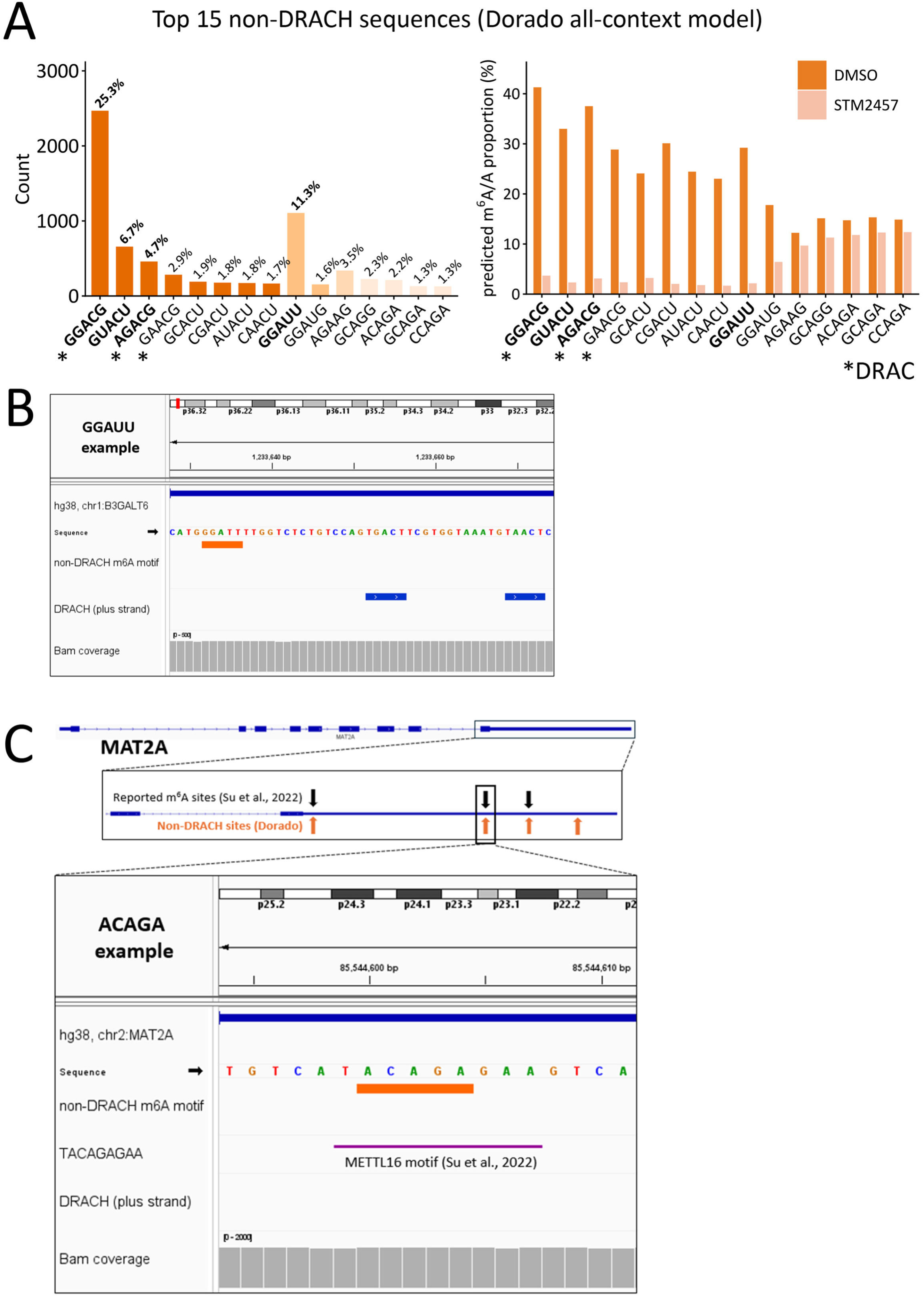
Non-DRACH m^6^A calls predicted by the Dorado all-context model in NHDF data. **(A)** Same as in Fig. S23A, but for m**^6^**A sites predicted with the all-context Dorado model. **(B)** Representative locus showing an STM2457-sensitive GGAUU call without evidence of positional shift, sequence variation, or homopolymer context. **(C)** STM2457-insensitive ACAGA calls within *MAT2A*, shown relative to the METTL16 consensus (UACAGAGAA).

**Figure S25.**
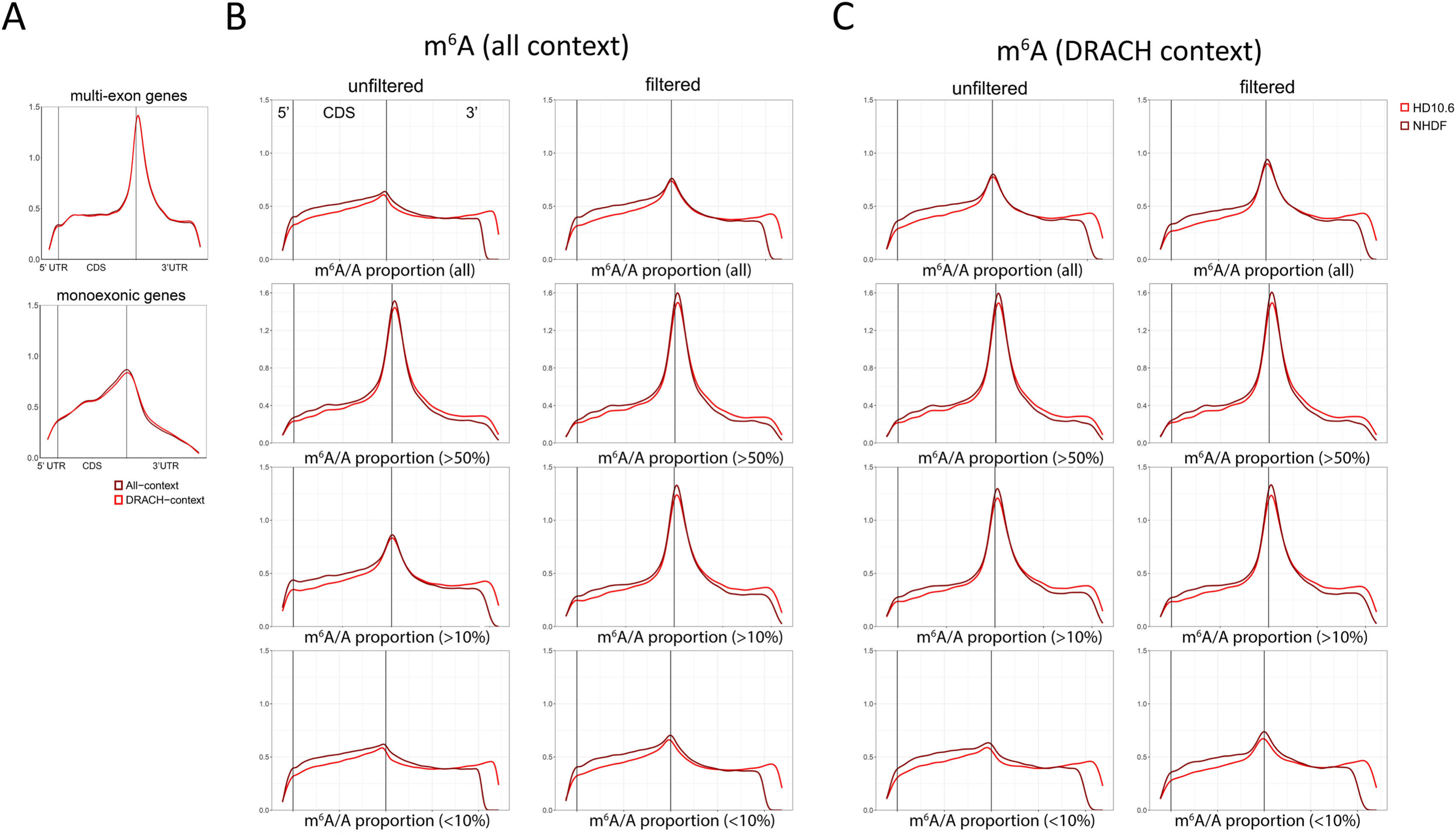
Metagene plots showing m^6^A (all-context and DRACH-context) distribution profiles in NHDF and HD10.6 data filtered at different predicted m^6^A/A proportions. **(A)** Metagene plots showing the density of m^6^A (all-context – dark red, DRACH-context – bright red) sites across all annotated poly(A) transcripts (upper plot) and monoexonic poly(A) transcripts (lower plot) from genome-aligned DMSO-treated NHDF datasets filtered with 0.98 modification probability, ≥ 20 coverage, and ≥ 10% predicted m^6^A/A proportion. **(B)** The density of all-context m^6^A sites (0.98 modification probability-unfiltered – left, filtered – right) is plotted across all annotated poly(A) transcripts of NHDF (dark red) and HD10.6 (bright red) datasets, filtered with different predicted m^6^A/A proportions (descending: all proportions, ≥ 50%, ≥ 10% and < 10%. All datasets were filtered additionally for ≥ 20 coverage. **(C)** Same as (A), but for DRACH-context m^6^A sites.

